# Faster and more accurate assessment of differential transcript expression with Gibbs sampling and edgeR v4

**DOI:** 10.1101/2024.06.25.600555

**Authors:** Pedro L. Baldoni, Lizhong Chen, Gordon K. Smyth

## Abstract

Differential transcript expression analysis of RNA-seq data is an increasingly popular tool to assess changes in expression of individual transcripts between biological conditions. Software designed for transcript-level differential expression analyses account for the uncertainty of transcript quantification, the read-to-transcript ambiguity (RTA), in statistical analyses via resampling methods. Bootstrap sampling is a popular resampling method that is implemented in the RNA-seq quantification tools kallisto and Salmon. However, bootstrapping is computationally intensive and provides replicate counts with low resolution when the number of sequence reads originating from a gene is low. For lowly expressed genes, bootstrap sampling results in noisy replicate counts for the associated transcripts, which in turn leads to non reproducible and unrealistically high RTA-dispersion for those transcripts. Gibbs sampling is a more efficient and high resolution algorithm implemented in Salmon. Here we leverage the developments of edgeR v4 to present an improved differential transcript expression analysis pipeline with Salmon’s Gibbs sampling algorithm. The new bias-corrected quasi-likelihood method with adjusted deviances for small counts from edgeR, combined with the efficient Gibbs sampling algorithm from Salmon, provides faster and more accurate DTE analyses of RNA-seq data. Comprehensive simulations and test data show that the presented analysis pipeline is more powerful and efficient than previous differential transcript expression pipelines while providing correct control of the false discovery rate.

## Introduction

RNA sequencing (RNA-seq) is a powerful technology used in biomedical research to profile the transcriptome of living organisms [1]. Analysis of RNA-seq data often involves the assessment of differential expression of genomic features between biological conditions of interest, such as treated vs control samples, tumor vs healthy tissues, or genetically modified vs wildtype organisms [2]. Throughout the years, differential expression analysis of RNA-seq data has often been conducted with a gene-centric view [3]. In this context, genomic features are represented by genes, exons, or exon-exon junctions, and analyses are performed to detect gene-level differential expression or differential splicing between conditions [4, 5, 6, 7]. Meanwhile, an alternative approach is to focus directly on transcript isoforms and to evaluate differential expression for each specific isoform (or transcript) [8, 9]. Recently, the development of lightweight quantification tools *kallisto* [10] and *Salmon* [11] has reduced the computational cost of isoform-centric analyses. The new tools, together with reductions in short read sequencing costs, now make it practical to perform differential expression analyses of RNA-seq data at scale at the transcript level.

The *kallisto* and *Salmon* tools assume a fully annotated reference transcriptome to perform pseudo- and quasi-alignment of RNA-seq reads, respectively. Such a process entails creating a reduced representation of the sequencing data by unambiguously assigning sequence reads to equivalence classes (ECs) of transcripts. *Salmon* further accounts for sequencing biases and the distribution of fragment lengths to subdivide ECs into subclasses for improved quantification [12]. Because sequence reads are often compatible with multiple annotated transcripts, a phenomenon that we call *read-to-transcript ambiguity* (RTA) [13], individual transcript abundances are then estimated probabilistically from EC counts. While *kallisto* and *Salmon* both implement a standard expectation-maximization (EM) algorithm to obtain maximum likelihood estimates for the number of sequence reads deriving from each annotated transcript, *Salmon* further implements a variational Bayes EM (VBEM) as an alternative optimization algorithm. The use of pseudo- or quasi-alignment makes *kallisto* and *Salmon* computationally faster than any splice-aware genome aligner designed for short read sequencing data while providing measurements of the relative expression of individual transcripts directly.

The effect of RTA on transcript quantification precision can be estimated via resampling techniques [13]. One such technique that is implemented in both *kallisto* and *Salmon* is bootstrap sampling. Bootstrap sampling resamples from the RNA-seq sequence reads for each sample, mimicking the effect of technical replicates arising from resequencing, and repeats the transcript quantifications for each resample. More precisely, parametric bootstrapping is performed with a multinomial model for ECs with probabilities computed from observed EC counts, a process that is equivalent to but more efficient than an ordinary bootstrap applied to the original sequence reads. The number of sequence reads deriving from each annotated transcript is estimated from each bootstrap sample with either EM or VBEM algorithms. The number of bootstrap samples to be generated is specified by the user. Quantifying a single RNA-seq library with either *kallisto* or *Salmon* will result in the specified number of bootstrap replicate counts to be output for every annotated transcript.

We showed recently that RTA produces overdispersed transcript counts and that the amount of overdispersion can be estimated with a high degree of accuracy by fitting a quasi-Poisson model to the bootstrap samples for each transcript [13]. We further showed that the RTA-induced dispersion can be (literally) divided out of the transcript counts, after which the divided counts can be input into standard *edgeR* analysis pipelines that were originally developed for gene-level counts. The RTA-dispersions interfere with the RNA-seq mean-variance relationship if not accounted for but, after being divided out, the scaled counts show the same smooth mean-variance relationships that are familiar from gene-level analyses. The approach is implemented in the *catchSalmon* and *catchKallisto* functions of the *edgeR* package. These functions read the *kallisto* or *Salmon* output, including bootstrap samples, and estimate the RTA-dispersions. The *edgeR* divided-count analysis approach was shown to outperform previous differential transcript expression (DTE) pipelines in terms of statistical power, false discovery rate (FDR), speed and flexibility [13]. Our study assumed 100 bootstrap resamples per RNA-seq library and was conducted with *edgeR* v3 rather than the new *edgeR* v4 [14].

Despite the success of the *edgeR* divided-count approach, there are some disadvantages to relying on bootstrap samples. The first is that bootstrap resampling adds considerably to computational time and therefore dissipates one of the key advantages of the lightweight alignment tools. Bootstrap sampling is by far the most computationally demanding step in the analysis pipeline, consuming much more time than either the alignment or the statistical analysis.

Another disadvantage of bootstrapping is potential loss of resolution for genes with low coverage or for genes where all the sequence reads are assigned by chance to a single EC. Bootstrapping is, by definition, limited to repetitions of sequence reads that were observed. If all the reads for a particular multi-transcript gene are assigned to a single EC, then that will necessarily also be true for all reads from all the bootstrap samples as well. In this case, RTA overdispersion will be judged to be absent, perhaps unrealistically so. Another scenario occurs when a multi-transcript gene is lowly expressed but the reads are spread over different ECs. In this case, bootstrap sampling may result in highly variable replicate counts for the transcripts of that gene and to poorly estimated RTA-dispersion parameters because of highly granular resampling of the reads.

*Salmon* implements the uncollapsed Gibbs sampling algorithm from Turro *et al*. [15] as an alternative to bootstrap resampling. Gibbs sampling involves drawing transcript counts from a posterior multinomial model (conditional on estimated transcript abundances) followed by drawing transcript abundances from a posterior gamma model (conditional on estimated transcript counts). Gibbs sampling coupled with VBEM optimization (the default in *Salmon*) employs a Bayesian framework with strong priors. The Bayesian framework gives non-zero probability to unobserved ECs and is able to obtain higher resolution resampling counts that are not limited to single EC scenarios or to granular resampling of reads in low count situations. Moreover, Gibbs sampling results in transcript replicate counts directly and does not require model optimization during every iteration, making it much faster than bootstrap sampling.

Another area for possible improvement is the statistical treatment of small counts. Transcript-level analyses tend to generate smaller counts than gene-level analyses because of the larger number of transcripts compared to genes. *edgeR* v4 was released recently and features an improved quasi-likelihood (QL) treatment of small counts. It implements bias-adjusted deviances that allow unbiased estimation of quasi-dispersions even for genes or transcripts at very low expression levels. *edgeR* v4 moreover allows much faster analysis of large datasets because of the possibility of bypassing the estimation of transcript-wise negative binomial (NB) dispersions in favour of greater reliance on quasi-dispersions [14].

The purpose of the current article is three-fold. First, we compare bootstrap with Gibbs sampling for evaluating DTE. We find that Gibbs sampling not only decreases computation time but also improves statistical power and increases the number of transcripts than can be retained in the DTE analysis. Second, we replace *edgeR* v3 with *edgeR* v4. Again, we find that *edgeR* v4 not only decreases computational time but also improves statistical power and increases the number of transcripts than can be retained in the analysis. Third, we consider whether the number of technical samples can be reduced without materially affecting the results. We find that only a couple of hundred technical replicates are required in total, across all samples in the study, dramatically reducing the number of resamples that are recommended per library for large studies. Together, these changes result in significant improvements to an already very competitive DTE pipeline. In particular, all three refinements improve the speed of the pipeline, especially for datasets with a large number of samples.

As in our earlier study, the *edgeR* divided-count strategy is compared with *Swish* and *sleuth. Swish* [16] uses Wilcoxon tests over replicate samples that are averaged to assess DTE between two conditions of interest, with independent or paired samples, for single or multi-batch RNA-seq experiments via stratification. *Swish* adjusts transcript replicate counts for sample-specific transcript length and sequencing depth via median-ratio size factors. *sleuth* [17] tests for DTE under a linear measurement error model that decomposes the total variance into a biological variance component and an inferential variance component resulting from RTA that is estimated from transcript replicate counts. *sleuth* normalizes counts using sample-specific median-ratio size factors [18] followed by a started log-transformation to ensure positivity and normality. While both *edgeR* and *Swish* accept as input RNA-seq quantification from both *kallisto* and *Salmon*, with either bootstrap or Gibbs sampling, *sleuth* accepts RNA-seq quantification output only from *kallisto*.

The improved DTE analysis pipeline presented here leverages the highly streamlined bias-corrected QL pipeline from *edgeR* v4 and the efficient Gibbs sampling algorithm from *Salmon*. The quasi-Poisson model from Baldoni *et al*. [13] is fitted to transcript counts obtained from Gibbs sampling without further modification. Transcript-specific RTA-dispersion parameters are estimated and used to scale counts so that their effective size reflects their true precision. The latest QL pipeline with adjusted deviances for small counts from *edgeR* v4 bypasses the estimation of transcript-specific NB-dispersions and DTE can be assessed via quasi-F-tests [14].

The *edgeR* divided-count pipeline is implemented in the *edgeR* package [14] available from Bioconductor [19]. The *edgeR* function *catchSalmon* imports transcript counts and associated Gibbs resamples from *Salmon* to estimate the transcript-specific RTA-dispersions. Downstream DTE analyses can then be conducted on scaled transcript counts using the bias-corrected QL pipeline with adjusted deviances with the function *glmQLFit* as for gene-level analyses [14]. DTE analysis with Gibbs sampling is shown to be more powerful and faster than using bootstrap sampling while correctly controlling the false discovery rate using simulations designed with the Bioconductor package *Rsubread* [20]. The improvements of the presented DTE pipeline are further demonstrated on a case study of short read RNA-seq data from human lung adenocarcinoma cell lines.

## Materials and methods

### Simulated datasets

#### Simulation of RNA-seq sequence reads

We generated RNA-seq experiments with using the simulation framework developed by Baldoni *et al*. [13]. Briefly, libraries with 100bp paired-end sequence reads were simulated in FASTQ file format with the Bioconductor package *Rsubread* from a set of 41,372 protein-coding and lncRNA expressed reference transcripts of the mouse Gencode transcriptome annotation M27 (NCBI Gene Expression Omnibus series GSE60450). RNA-seq experiments from two groups were simulated in scenarios with either three, five, or ten biological replicate samples per group. Following Law *et al*. [6] and Baldoni *et al*. [13], transcript expression levels were generated to have a typical biological coefficient of variation (BCV) between replicates of 0.2 with higher BCVs for lower expression levels and inter-transcript BCV heterogeneity defined by prior degrees of freedom of 40. This matches the biological variability and heterogeneity that we typically observe our own in-house RNA-seq experiments using genetically identical laboratory mice [6]. Experiments were simulated with alternating library sizes of 25 Mi. or 100 Mi. paired-end reads over samples. For each scenario, 20 simulated experiments were generated. A random subset of 3,000 transcripts had their baseline relative expression adjusted with a two-fold-change to establish differential expression between groups with up- and down-regulated transcripts. For every scenario, simulations without any real differential expression between groups (null simulations) were also generated to assess methods’ type I error rate control.

A second set of simulations was conducted to emulate large-scale observational RNA-seq studies with human subjects, with 100 samples per group and with BCV increased to 0.6 [6].

#### Quantification of RNA-seq experiments

Simulated RNA-seq experiments were quantified with *Salmon* and *kallisto* with 100 bootstrap resamples per library. Experiments were re-quantified with *Salmon* and Gibbs sampling to generate 100 Gibbs resamples per library. Other options in *Salmon* and *kallisto* were left as default. Simulated RNA-seq reads were quantified with *Salmon* and *kallisto* with index generated from the complete mouse Gencode transcriptome annotation M27. For *Salmon*, we used a decoy-aware mapping-based indexed transcriptome generated from the mouse mm39 reference genome with k-mers of length 31. The R package *wasabi* (https://github.com/COMBINE-lab/wasabi) was used to convert *Salmon*’s output to *abundance*.*h5* files for DTE analyses in *sleuth*. Transcript quantification was performed in an Intel(R) Xeon(R) Gold 6342 (2.80GHz) processor with 10 threads used concurrently.

#### Assessment of differential transcript expression

We assessed the performance of *edgeR* with count scaling and other popular methods with respect to power to detect DTE, false discovery rate (FDR) control, type I error rate control, and computational speed. Two versions of *edgeR* with scaled counts were evaluated, herein denoted as *edgeR*-v3 and *edgeR*-v4. For *edgeR*-v3, we used the previous QL pipeline with the weighted likelihood empirical Bayes method implemented in *estimateDisp* to obtain posterior NB-dispersion estimates [21, 13]. For *edgeR*-v4, we used the QL pipeline with common NB-dispersion estimated using the top 5% of most highly expressed genes and quasi-dispersions estimated from the bias-adjusted deviances [14]. Other methods benchmarked in our study were *sleuth* with likelihood ratio test (*sleuth-LRT*), *sleuth* with Wald test (*sleuth-Wald*), and *Swish* (implemented in the Bioconductor package *fishpond*). Analyses were performed with transcript resamples generated with either the bootstrap or Gibbs sampling methods. For *edgeR*, low-expression transcripts were filtered by *filterByExpr*, library sizes were normalized by *normLibSizes*, and differential expression was assessed by quasi-F-tests with default options. For *sleuth* and *Swish*, default filtering and pipeline options implemented in their respective packages were used throughout our simulations. In all analyses, transcripts were considered to be differentially expressed (DE) under an FDR control of 0.05.

### Human lung adenocarcinoma cell lines

Illumina short-read paired-end RNA-seq libraries were obtained from NCBI Gene Expression Omnibus (GEO) series GSE172421. Three biological replicate samples were used to examine the transcriptomic profile of human adenocarcinoma cell lines NCI-H1975 and HCC827. RNA-seq samples were quantified in *Salmon* with bootstrap and Gibbs sampling algorithms using 10 threads concurrently. A total of 100 bootstrap and Gibbs resamples were generated for every RNA-seq library. Other options in *Salmon* were left as default. A decoy-aware mapping-based indexed transcriptome generated from the human hg38 reference genome with k-mers of length 31 and the complete human Gencode transcriptome annotation version 33 was used during quantification. The *edgeR* function *catchSalmon* was used to import *Salmon*’s quantification and estimate the RTA-dispersion parameter with either bootstrap or Gibbs sampling. Transcript counts were scaled, non-expressed transcripts were filtered by *filterByExpr*, genes other than protein-coding and lncRNA were removed, and library sizes were normalized by *normLibSizes*. Differential expression was assessed with quasi-F-tests under both *edgeR*-v3 and *edgeR*-v4 pipelines [13, 14]. Sequence reads from the RNA-seq experiments were aligned to the hg38 reference genome with the Bioconductor package *Rsubread* for assessment of read coverage.

### Dividing out the read-to-transcript ambiguity

The distributional assumptions made by *edgeR* have been described by Chen *et al*. [14]. We summarize the assumptions here with some notational changes to emphasize the role of the RTA in the context of DTE. Write *y*_*ti*_ for the fractional number of reads by assigned by *Salmon* or *kallisto* to transcript *t* in sample *i*. The systematic part of *y*_*ti*_, *µ*_*ti*_ = *E*(*y*_*ti*_), can be modelled by a log-linear model

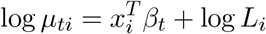

where *x*_*i*_ is a covariate vector specifying the experimental conditions applied to sample *i, β*_*t*_ is a coefficient vector that captures the experimental effects and log-fold-changes, and *L*_*i*_ is the effective library size for sample *i*.

*edgeR* assumes that technical replicates produced from the same RNA sample (and with the same library size) would result in quasi-Poisson repeatability with variance 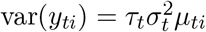, where *τ*_*t*_ is the dispersion arising from RTA and 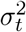 is the dispersion arising from other technical factors such as PCR duplication. We assume that *τ*_*t*_ ≥ 1 and 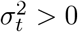

*edgeR* further assumes that the true expression level of transcript *t* varies between biological replicates with squared coefficient of variation equal to *ψ*_*t*_. The read count distribution therefore follows a mixture distribution across biological replicates with a quadratic mean-variance relationship of the form

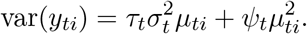

The RTA dispersion *τ*_*t*_ depends on the annotation topology of the gene that the transcript belongs to whereas 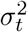 and *ψ*_*t*_ depend on the biological samples and on the sequencing technology. The *τ*_*t*_ can be obtained via resampling techniques but 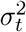 and *ψ*_*t*_ must be estimated from the biological replicates using linear modelling. The 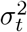 and *ψ*_*t*_ parameters tend to be abundance-dependent between transcripts whereas the *τ*_*t*_ are far more transcript-specific and vary by several orders of magnitude between transcripts.

Baldoni *et al*. [13] presented a method for estimating the RTA-dispersions *τ*_*t*_ from the bootstrap or Gibbs resamples generated by *Salmon* or *kallisto*. The estimate 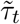 is equal to the squared coefficient of variation of the resampled transcript counts, bounded below by 1, and with very light empirical Bayes moderation between transcripts to add stability when the counts are very small. The estimates are averaged over all samples in the study, so the degrees of freedom for estimating *τ*_*t*_ is equal to *N* (*B* − 1) where *N* is the total number of RNA samples and *B* is the number of *in silico* resamples for each RNA sample. If *N* (*B* −1) is large, then the estimate of *τ*_*t*_ will be very precise.

Baldoni *et al*. also showed that RTA dispersions can be divided out of the transcript read counts to obtain scaled counts with a more standard mean-variance relationship. Define the divided counts by 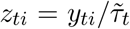. If 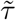 is a precise estimate of *τ*, then

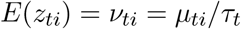

and

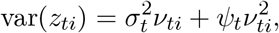

which can alternatively be written as

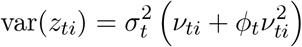

where 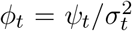. This is the standard variance function for a quasi-negative-binomial generalized linear model, with *ϕ*_*t*_ as the NB-dispersion parameter and 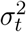 as the quasi-dispersion. The divided counts can therefore be analyzed directly using the *edgeR* QL workflow originally developed for gene-level differential expression analyses. The fact that the RTA dispersions have been removed ensures that the data should behave similarly to gene-level counts and the analysis should proceed with full statistical efficiency.

Note that the divided count analysis pipeline outlined above is very robust to under-estimation or over-estimation of the RTA dispersions, because any consistent multiplicative bias in 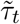 will be absorbed into the quasi-dispersions 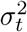. On the other hand, the 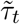 estimates should be proportional to the true *τ*_*t*_ in order that the variance of the divided counts be independent of transcript annotation topology.

### Bias-corrected quasi-likelihood method

In practice, it isn’t possible to estimate transcript-specific values for both 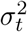 and *ϕ*_*t*_, so *ϕ*_*t*_ is set to a global value and transcript-specific variation is captured by 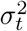. The 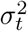 are estimated by trended empirical Bayes from the generalized linear model residual deviances [22, 23]. *edgeR*-v3 uses trended NB binomial dispersions [21] for *ϕ*_*t*_ whereas *edgeR*-v4 sets all the *ψ*_*t*_ to a common value estimated from the top most highly expressed genes.

A review of *edgeR* v4 is given by Chen *et al*. [14]. For transcripts with very low counts, the traditional chisquare approximation to the NB deviances is biased and results in underestimation of the quasi-dispersions. *edgeR*-v4 instead implements adjusted NB deviances with adjusted degrees of freedom so that the chi-square approximation is exactly correct for the first two moments (mean and variance). As a result, the new method provides unbiased estimation of the quasi-dispersions and improved statistical power in DTE analyses. The accuracy of the new QL method allows the biological coefficient of variation (BCV) to be set to a common value for all transcripts (estimated from the top most highly expressed transcripts), bypassing the computationally intensive estimation of individual NB-dispersions via *estimateDisp* and improving on computational speed from previous QL pipelines.

### Usage and implementation

The calculation of RTA-dispersions 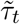 is implemented in the *edgeR* functions *catchSalmon* and *catchKallisto*. The functions return the matrix of transcript counts together with the associated RTA-dispersions. *catchSalmon* automatically detects and accepts resampled transcript counts from either Gibbs or bootstrap sampling algorithms. Divided counts *z*_*ti*_ are computed from the *catchSalmon* or *catchKallisto* output and then enter the *edgeR* pipelines as regular read counts. *edgeR* accepts fractional counts directly, so the divided counts do not need to be rounded to integers [13, 14].

The bias-corrected QL method with adjusted deviances is implemented in the *edgeR* function *glmQLFit*. For backward compatibility, *glmQLFit* allows users to switch between the traditional and adjusted deviances by setting *legacy=TRUE* or *legacy=FALSE*. The legacy QL pipeline requires trended NB-dispersions to be estimated by *estimateDisp* before running *glmQLFit*, but the new pipeline estimates a common BCV internally from the most highly expressed transcripts. In this article, all analyses were conducted by *edgeR* 4.0.16. Analyses using *legacy=TRUE* are denoted as *edgeR*-v3 while the new pipeline with *legacy=FALSE* is denoted as *edgeR*-v4.

## Results

### Gibbs and bootstrap sampling yield correlated RTA-dispersions but with systematic differences

We first explored the differences between the RTA-dispersion estimates obtained with either Gibbs or bootstrap sampling from *Salmon* for the lung adenocarcinoma cell line data. We observed that the Gibbs and bootstrap RTA-dispersions were highly correlated but that the bootstrap values were on average higher (Figure 1a). For example, the maximum bootstrap RTA-dispersion was about 2^10^ compared to about 2^8^for Gibbs sampling. On the other hand, there were a number of transcripts seen on the far left of Figure 1a that have no overdispersion assessed by bootstrap but have dispersion as high as 2^5^by Gibbs sampling. Figure 1b shows that bootstrap and Gibbs RTA-dispersions both tend to decrease with count size. The bootstrap vs Gibbs difference also decreases with count size, with bootstrap RTA-dispersions markedly larger than Gibbs RTA-dispersions for log2-CPM*<* 0 but almost equal on average for log2-CPM*>* 2. The fact that Gibbs sampling tends to return lower RTA-dispersions than the bootstrap for low-count transcripts is presumably due to the strong Bayesian prior employed by *Salmon* in its Gibbs sampling algorithm, and this is explored by way of examples in the next Section.

**Figure 1.**
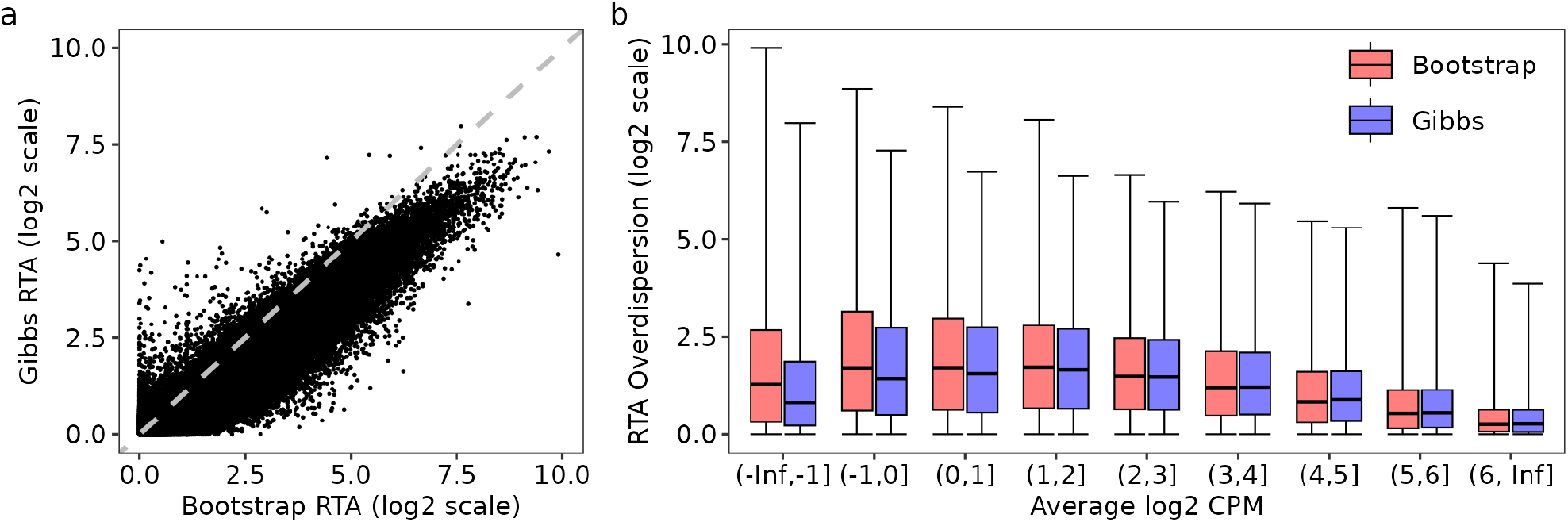
Gibbs vs bootstrap transcript-level RTA-dispersion estimates from *Salmon* for the lung adenocarcinoma cell line data. (a) scatterplot of Gibbs vs bootstrap-based RTA-dispersion estimates (log2 scale). (b) boxplot showing Gibbs and bootstrap-based RTA-dispersions as a function of the average log2 CPM of each transcript. In panels (a) and (b), RTA-dispersion estimates were obtained with 100 resamples. In panel (a), the line of equality is shown as a gray dashed line. In panel (b), average log2 CPM values were computed from scaled counts obtained with Gibbs RTA-dispersion estimates. All 137,788 expressed transcripts are shown.

RTA-dispersions estimated from *kallisto* bootstrap samples were highly correlated with those from *Salmon*, and tended to be intermediate between the *Salmon*-bootstrap and *Salmon*-Gibbs RTA-dispersions (Supplementary Figure S1).

### Gibbs sampling has better resolution than bootstrap sampling

Bootstrap sampling can be problematic in specific scenarios in which Gibbs sampling can provide more precise RTA-dispersion estimates and, consequently, more accurate DTE assessment. The first scenario is when a shared EC is dominated by a single transcript that receives all sequence reads from that EC during probabilistic assignment. Bootstrap sampling will result in no variation in transcript assignment for those reads, resulting in simple multinomial variation over the technical replicates for the dominant transcript and zero counts for other transcripts. This scenario often occurs when annotated transcripts differ by a single exon and no sequence read is compatible with that particular exon. In the lung adenocarcinoma cell lines data, this phenomenon was observed for transcripts *HABP2-201* and *HABP2-203* of the tumor suppressor gene *HABP2* [24]. Bootstrap sampling resulted in an estimated RTA-dispersion of 1.00 (no overdispersion) for *HABP2-201* and *HABP2-203*, an unrealistic RTA measurement for the short dominating transcript *HABP2-201* (Figure 2a). Effective counts computed via count scaling under Gibbs sampling were approximately 30% smaller than those computed via bootstrap sampling for transcript *HABP2-201*, which likely better reflects the precision associated with RTA introduced during transcript quantification.

**Figure 2.**
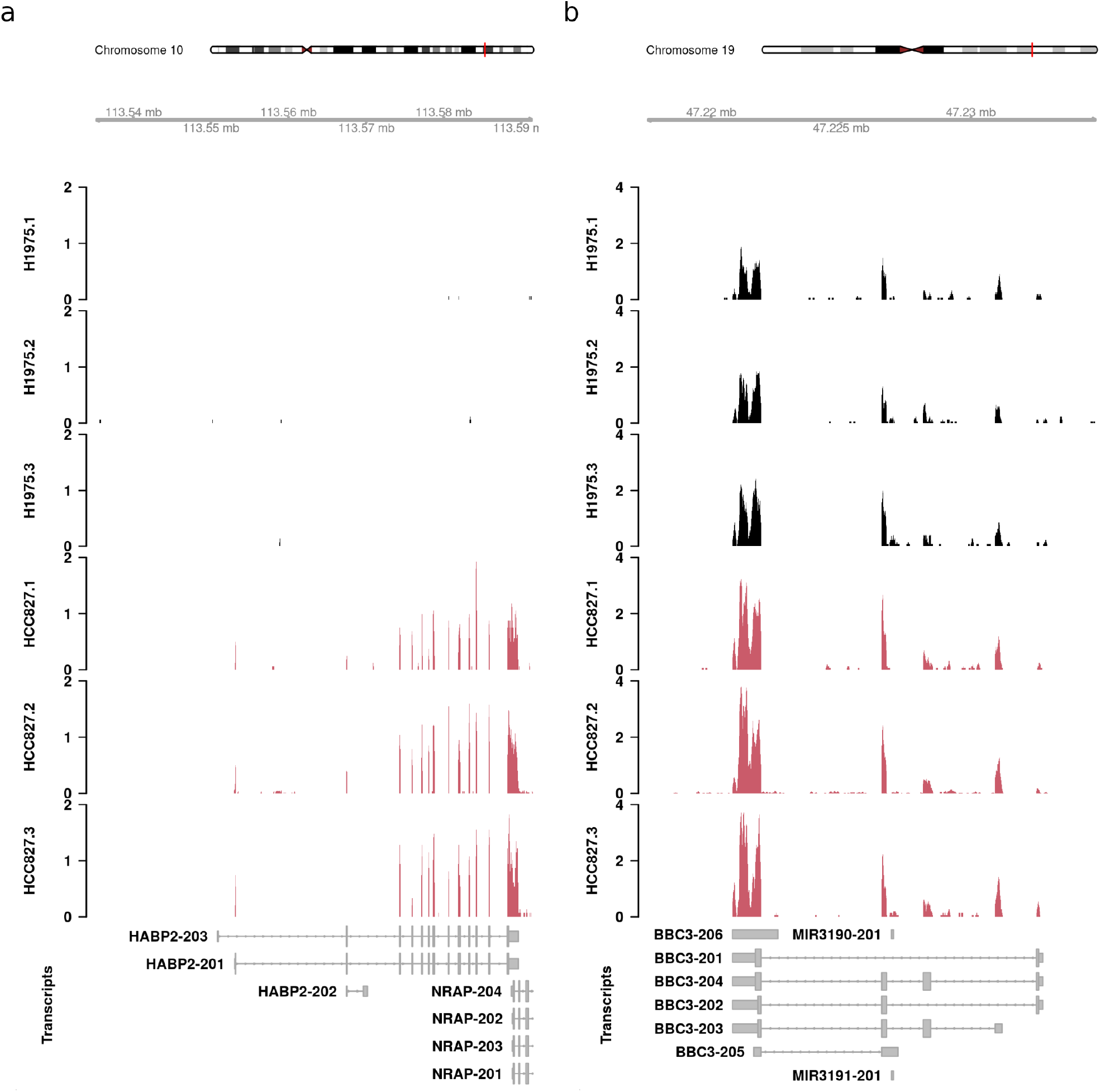
Coverage tracks from the human lung adenocarcinoma cell lines data. In panel (a), coverage profile of the cancer gene *HABP2*. In every HCC827 sample, all ambiguous reads are probabilistically assigned to transcript *HABP2-201*, which results in no overdispersion under bootstrap sampling with an estimated RTA-dispersion of 1.00. The estimated RTA-dispersion under Gibbs sampling was 1.42. In panel (b), coverage profile of the pro-apoptotic gene *BBC3*. The lowly abundant transcript *BBC3-202* (average log CPM -0.433) is filtered out upon count scaling with bootstrap sampling but reported as DE with Gibbs sampling (1.35 log fold-change, adjusted p-value 0.006). Coverage profiles are defined as the number of reads overlapping each base, transformed into a count-per-million value based on library size. Coverage tracks include all protein-coding and lncRNA transcripts based on the human Gencode annotation version 33.

A second and more common scenario is when a multi-transcript gene is lowly expressed. In this case, bootstrap sampling will result in noisy replicate counts for each of the annotated transcripts of that gene. As a result, bootstrap sampling will fail to accurately reproduce the variability associated with resequencing. This phenomenon leads to an increased RTA-dispersion for lowly expressed transcripts which, in turn, may cause loss of power in statistical analyses. An interesting example was the *BBC3-202* transcript from the pro-apoptotic gene *BBC3* [25]. Transcript *BBC3-202* was found to be DE between NCI-H1975 and HCC827 cell lines under Gibbs sampling but excluded from the analysis upon count scaling with bootstrap sampling (Figure 2b). We observed 1,303 similar cases in the analysis of the lung adenocarcinoma cell lines data (Supplementary Figure S2).

### 0.1 edgeR accommodates systematic differences between resampling algorithms

As noted in Methods, systematically lower overdispersion estimates from Gibbs as compared to bootstrap resampling will not generally impact the FDR or statistical power achieved by *edgeR* because *edgeR*’s QL pipeline will correct any consistent bias in the RTA-dispersions according to the biological variability observed between replicates in the downstream DTE analysis. The effectiveness of the RTA-dispersion estimates in the *edgeR* pipeline will be determined more by their correlation with the true RTA-dispersions than by their absolute size. Visually, dividing by Gibbs RTA-dispersions was found to stabilize the estimation of NB-dispersions in *edgeR* just as well as dividing by the bootstrap versions (Supplementary Figure S3).

On the other hand, lower RTA-dispersion estimates will result in divided-counts that are not as small and hence will increase the number of transcripts included in the downstream analysis after filtering.

### Gibbs sampling and edgeR v4 provide more accurate assessment of differential transcript expression

We used the simulation framework from Baldoni *et al*. [13] to assess differences between bootstrap and Gibbs sampling in regard to DTE assessment. We benchmarked both resampling algorithms with respect to downstream metrics from DTE methods such as power to detect DE transcripts, ability to control the FDR, and type 1 error rate control. Our analyses included *edgeR* with count scaling under both v3 and v4 pipelines as well as other previously benchmarked DTE methods reported in [13]. Figure 3 presents the main results from our simulations with five samples per group.

**Figure 3.**
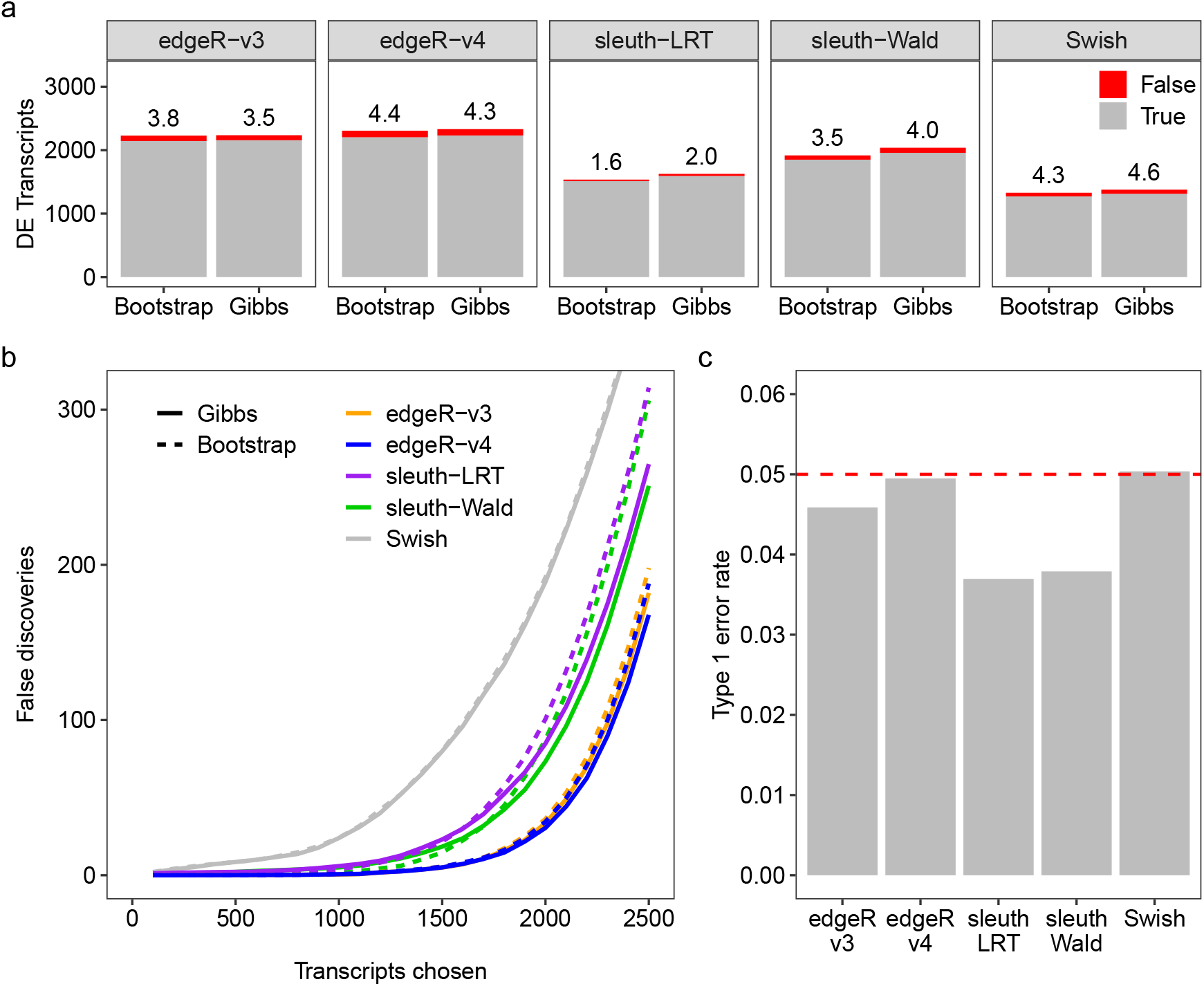
Power and error rate control for simulated data with 100bp paired-end reads, unbalanced library sizes, and five samples per group. **(a)** stacked barplots show the number of true (gray) and false (red) positive DE transcripts at nominal 5% FDR for different DTE and sampling methods. Observed FDR is shown as a percentage over each bar. **(b)** number of false discoveries as a function of the number of transcripts chosen for different DTE and sampling methods. **(c)** observed type 1 error rate for the different DTE methods calculated as the proportion of transcripts with unadjusted p-values *<* 0.05 in null simulations (without differential expression) quantified with Gibbs sampling. Dashed line indicates the expected proportion of p-values *<* 0.05 under the null hypothesis of no differential expression. Transcript quantification was performed with *Salmon* with 100 bootstrap or Gibbs resamples. Results are averaged over 20 simulations.

In the *n* = 5 scenario considered by Figure 3, all the DTE methods were found to control the error rate correctly either in terms of FDR (Figure 3a) or in terms of type 1 error rate when no truly DE genes were present (Figure 3c). Figure 3b shows the operating characteristics of each DTE method in terms of false discoveries versus number of transcripts with transcripts ranked by p-value. This shows *edgeR*-v4 to have the lowest number of false discoveries at any chosen number, followed by *edgeR*-v3, *sleuth-Wald, Sleuth-LRT* and *Swish*. Except for *Swish*, all the methods were improved by substituting Gibbs in place of bootstrap sampling.

Gibbs sampling led to detection of more truly DE transcripts than bootstrap sampling for all benchmarked methods at the 0.05 FDR level (Figure 3a). The absolute (relative) increase in the number of truly DE transcripts detected by each method was 108 (6%), 82 (5%), 42 (3%), 28 (1%), and 12 (1%) for *sleuth-LRT, sleuth-Wald, Swish, edgeR-v4* and *edgeR-v3* respectively, averaged over simulations. *edgeR*-v4 was the most powerful DTE method, detecting the greatest number of truly DE genes with either resampling method (Figure 3a). On average, *edgeR*-v4 detected 60 to 120 more truly DE transcripts than its previous version.

Simulations were also conducted with *n* = 3, *n* = 10 or *n* = 100 samples per group (Supplementary Figures S4, S5 and S11). The results were qualitatively similar to those for *n* = 5 in that *edgeR*-v4 was consistently the most accurate analysis pipeline, followed by *edgeR*-v3, and Gibbs sampling consistently gave slightly better results than bootstrap sampling. However, *Swish* failed to control the FDR correctly for *n* = 3 and *n* = 100 while *Sleuth* failed to control the FDR for *n* = 100. *Sleuth* also lacked power to detect any DE at the FDR= 0.05 level when *n* = 3.

Supplementary Figures S6–S8 present the corresponding plots using *kallisto*, with similar results to those obtained with bootstrapping from *Salmon*.

In summary, the simulations demonstrate the improvements in statistical power and accuracy provided by Gibbs sampling and *edgeR*-v4 either together or separately. *edgeR*-v3 and *edgeR*-v4 were the only analysis pipelines to correctly the FDR in all scenarios, and they also detected more truly DE transcripts than the output pipelines. In combination, *Salmon* and Gibbs sampling followed by *edgeR*-v4 analysis was the most powerful and accurate pipeline for DTE analysis.

### Gibbs sampling provides a faster quantification pipeline for differential transcript expression assessment

We evaluated the computational performance of bootstrap and Gibbs sampling algorithms during transcript quantification with *Salmon* using our simulated RNA-seq experiments. Resampling methods were benchmarked with respect to computing time when running *Salmon* with 10 threads and with the default VBEM algorithm during the offline optimization phase. Our results are summarized in Table 1.

**Table 1.** Total processing time per library with either *Salmon* bootstrap or Gibbs resampling method. The total processing time is further split into the time spent during quantification, including algorithm initialization and optimization steps, and the time spent to generate the 100 bootstrap or Gibbs resamples. Results are averaged over all paired-end simulated libraries.

| Resampling Method | Library Size<br>(million read pairs) | Quantification Time (minutes) |  |  |
| --- | --- | --- | --- | --- |
|  |  | Optimization | Resampling | Total |
| Bootstrap | 25 | 1.79 | 9.45 | 11.24 |
|  | 50 | 3.22 | 13.41 | 16.64 |
|  | 100 | 6.09 | 19.15 | 25.24 |
| Gibbs | 25 | 1.78 | 0.91 | 2.69 |
|  | 50 | 3.22 | 1.62 | 4.85 |
|  | 100 | 6.05 | 3.01 | 9.06 |

We observed that the total quantification time varied with respect to the library size and the choice of resampling method. With bootstrap sampling, the average total time varied between 11 and 25 minutes for libraries with 25 and 100 million reads pairs, respectively. In contrast, *Salmon* was substantially faster with Gibbs sampling with libraries being quantified in 3 and 9 minutes under equivalent library size configurations, on average. Differences in total quantification time between pipelines were a result of the choice of the sampling method alone for any given library size specification, as both pipelines used VBEM as the main optimization routine before resampling. The time spent by *Salmon* to draw bootstrap resamples represented 84% and 76% of the total quantification time for libraries with 25 and 100 million reads pairs, respectively. Conversely, Gibbs sampling was considerably faster and resulted in a resampling time that represented only 33% of the total quantification time regardless of the library size. For a typical RNA-seq library with 50 million read pairs, the average time for a single bootstrap resample to be drawn was approximately eight times the time for *Salmon* to draw a single resample using the Gibbs sampling algorithm. With bootstrap sampling, *kallisto* was approximately 45% faster than *Salmon*, taking between 5 and 12 minutes to quantify libraries with 25 and 100 million reads pairs, respectively (Supplementary Table S1). Yet, *Salmon* with Gibbs sampling ranks as the fastest quantification method available. For a typical RNA-seq library with 50 million read pairs and 30 resamples, our results suggest an approximate 50% reduction in total time for *Salmon* transcript quantification by using Gibbs sampling instead of bootstrapping.

### How many technical samples are required?

Reducing the number of bootstrap or Gibbs resamples during transcript quantification will obviously reduce compute time but may might also adversely affect the statistical performance of downstream DTE methods. The RTA-dispersions are estimated from a quasi-Poisson for which the residual degrees of freedom are equal to *N* (*B*− 1), where *N* is the total number of RNA-seq samples in the study and *B* is the number of technical replicates generated per sample. The residual degrees of freedom determine the precision with which the RTA-dispersions are estimated. Aiming to optimize computing time while maintaining full statistical efficiency, we evaluated the necessary number of resamples for DTE analyses. Specifically, we computed downstream observed power and FDR obtained with different number of Gibbs and bootstrap resamples. Our results using *edgeR*-v4 are presented in Figure 4 for three, five and ten samples per group and in Supplementary Figure S11 for 100 samples per group. Results using bootstrap samples from *kallisto* were similar to those with bootstrap samples from *Salmon* (Supplementary Figure S9).

**Figure 4.**
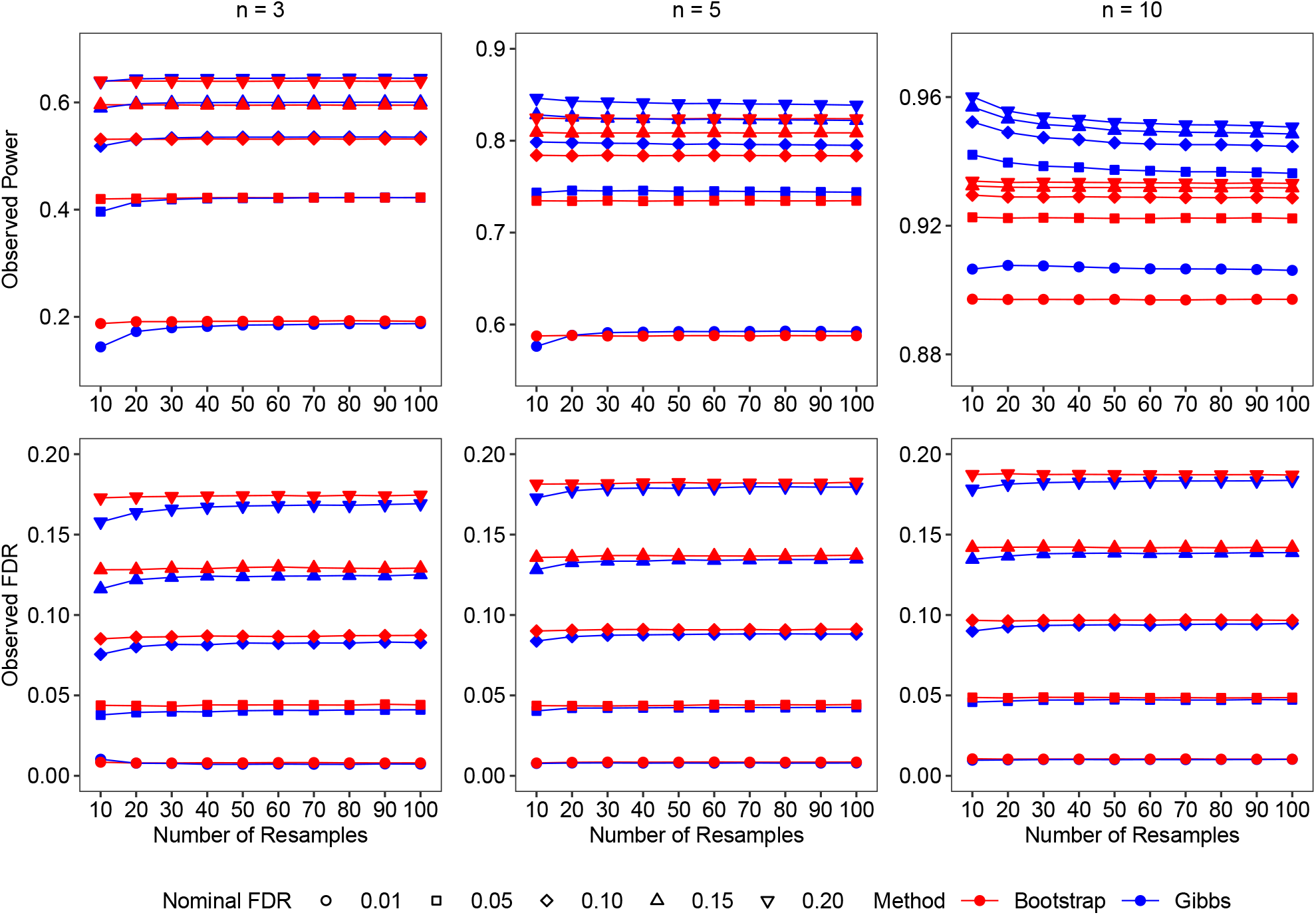
Plots showing the observed power and the observed false discovery rate (FDR) versus the number of replicate samples while controlling for different nominal FDR levels. Datasets were generated for two groups with *n* = 3, *n* = 5 or *n* = 10 samples per group. Results are presented for the different number of samples per group and for both Gibbs- and bootstrap-based DTE pipelines with *Salmon* and *edgeR*-v4. Simulated data generated with 100bp paired-end reads and unbalanced library sizes. Results are averaged over 20 simulations.

We note that *edgeR*-v4 correctly controlled the FDR below the nominal level for all sample sizes and for all FDR levels regardless of the number of resamples conducted or the type of resampling algorithm. Statistical power also was not very sensitive to the number of resamples conducted. When Gibbs samples were generated and a stringent FDR cutoff of 0.01 was used, *edgeR*-v4 reached maximum statistical power with *B* = 50 technical samples when *n* = 3, with *B* = 30 samples when *n* = 5, and with *B* = 20 when *n* = 10 (Figure 4). At less stringent FDR cutoffs, maximum power was reached even more quickly.

Since the total number of samples in the two groups is *N* = 2*n*, this suggests that Gibbs sampling can benefit from *N* (*B* − 1) up to about 200–300, although the increase in power was small and good results were already achieved with *N* (*B* − 1) *>* 60. For a large-sample experiment with *n* = 100 samples per group, full performance was achieved with just *B* = 2 Gibbs samples per RNA sample, corresponding to *N* (*B* − 1) = 200 (Supplementary Figure S11).

When bootstrap samples were generated, the results were even more stable with little change in statistical power with increasing number of technical samples. Close to maximum power was already achieved by *B* = 10 bootstrap samples for *n* = 3 and by *B* = 2 when *n* = 100 (Figure 4, Supplementary Figure S11).

### edgeR-v4 provides faster differential transcript expression analysis

Our simulated experiments compared the performance of the *edgeR*-v4 QL pipeline with that of *edgeR*-v3. Table 2 presents the analysis time consumed the two pipelines in our benchmarking study, and shows that *edgeR*-v4 enjoys a near 40% reduction in computing time. The faster computing time of *edgeR*-v4 over *edgeR*-v3 was a result of the newer estimation strategy of quasi-dispersions from *edgeR*, which allows the estimation of transcript-wise NB-dispersions to be completely bypassed. This has proven to be an effective strategy for the analysis of large datasets [14].

**Table 2.** Total analysis time (seconds), observed power, and observed false discovery rate (FDR) from *edgeR*-v3 and *edgeR*-v4 for different sampling methods and different number of samples per group. Nominal FDR of 0.05 in DE analyses. Analyses were performed with *Salmon* and 100 resamples. Results are averaged over 20 simulations.

| Samples per Group | Sampling Method | edgeR-v3 |  |  | edgeR-v4 |  |  |
| --- | --- | --- | --- | --- | --- | --- | --- |
|  |  | Time | Power | FDR | Time | Power | FDR |
| Three | Bootstrap | 6.79 | 0.39 | 0.04 | 3.54 | 0.42 | 0.04 |
|  | Gibbs | 7.81 | 0.38 | 0.03 | 4.46 | 0.42 | 0.04 |
| Five | Bootstrap | 9.96 | 0.71 | 0.04 | 5.79 | 0.73 | 0.04 |
|  | Gibbs | 11.38 | 0.72 | 0.04 | 7.21 | 0.74 | 0.04 |
| Ten | Bootstrap | 17.78 | 0.91 | 0.04 | 10.95 | 0.92 | 0.05 |
|  | Gibbs | 20.60 | 0.92 | 0.04 | 13.61 | 0.94 | 0.05 |

Gibbs sampling slightly increased the total analysis time of *edgeR* as a side effect of including more transcripts in the statistical analyses. The increase in analysis time was though more than compensated by the decrease in computing time required to generate Gibbs samples instead of bootstrapping.

Supplementary Figure S12 shows analysis times for all the DTE methods for the simulated data with *n* = 100 samples in each group. Again, *edgeR*-v4 is faster than *edgeR*-v3, and both are several times faster than *sleuth* or *Swish*.

### Differential transcript expression in human adenocarcinoma cell lines

Using the *edgeR*-v4 QL pipeline and *Salmon*’s Gibbs sampling algorithm, we performed a DTE analysis of the Illumina short paired-end read RNA-seq experiment from the human adenocarcinoma cell lines data. The RNA-seq experiment comprises 6 samples of NCI-H1975 and HCC827 cell lines with 3 biological replicate samples per cell line. Libraries were sequenced with an Illumina NextSeq 500 sequencing system, producing 28–134 million 80 bp read-pairs per sample. A previous DTE analysis of this dataset with *edgeR*-v3 and bootstrap sampling reported thousands of DE transcripts, many of which from known cancer-related alternatively spliced genes [13]. Gibbs sampling provides a faster and higher resolution alternative to bootstrapping during transcript quantification, and the v4 QL pipeline employs a bias-corrected QL method for more powerful DTE analyses. Hence, we expected to observe improvements in regard to statistical power and computing time in this analysis of the human adenocarcinoma cell lines data with *edgeR*-v4 and Gibbs sampling. In fact, switching from bootstrap to Gibbs sampling in *Salmon* resulted in a reduction of the quantification time from 22 minutes to 5 minutes per sample, on average. The new bias-corrected QL method with adjusted deviances for small counts resulted in quasi-dispersion estimates close to one even for transcripts with very small counts (Figure 5a).

**Figure 5.**
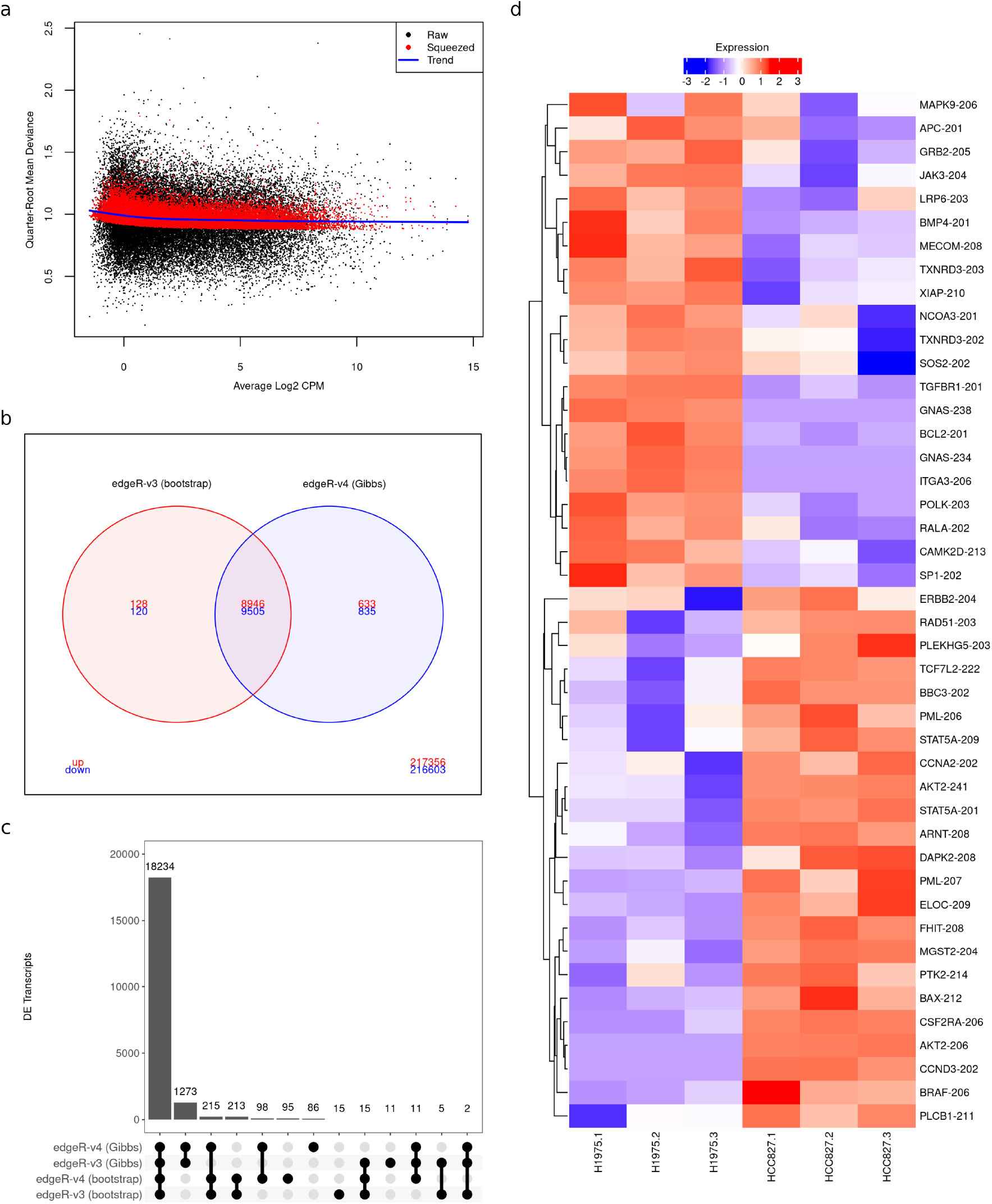
Panels (a)–(d) show the main results from the RNA-seq analysis of the human lung adenocarcinoma cell lines. In (a), quasi-dispersion plot showing the quarter-root of the quasi-dispersions from *edgeR*-v4 against the average log2 counts per million of each transcript. The plot shows the raw quasi-dispersions, equal to the mean residual deviance for each transcript, the trended empirical Bayes prior and the final (squeezed) posterior values. In (b), Venn diagram with DTE results from *edgeR*-v3 and *edgeR*-v4 with bootstrap and Gibbs sampling, respectively. The number of up- and down-regulated DE transcripts in the HCC827 cell line are indicated in red and blue, respectively. In (c), upset plot showing the number of DE transcripts from *edgeR*-v3 and *edgeR*-v4 with either bootstrap or Gibbs sampling. In (d), heatmap of transcripts associated with KEGG cancer and non-small cell lung cancer pathways found to be DE between NCI-H1975 and HCC827 cell line exclusively by the Gibbs DTE pipeline. RNA-seq libraries were quantified with *Salmon* with 100 resamples. Scaled log2 counts per million are displayed as expression levels in panels (c)–(d). Nominal FDR of 0.05 in transcript-level analyses with *edgeR*.

Our DTE analysis with Gibbs sampling and *edgeR*-v4 quasi-F-tests detected 19,919 DE transcripts between NCI-H1975 and HCC827 cell lines (9,579 up-regulated and 10,340 down-regulated transcripts in the HCC827 cell line; Figure 5b). The new analysis presented 1,220 additional DE transcripts than previously reported [13] and revealed 91 novel isoform-switching genes with up- and down-regulated DE transcripts.

The set of novel isoform-switching genes included the proto-oncogene *BRAF* (*BRAF-204*, -0.88 log foldchange, adjusted p-value 1.183 × 10^−5^; *BRAF-206*, 1.13 log fold-change, adjusted p-value 1.085 × 10^−2^) for which cancer-causing mutations in lung adenocarcinoma have been previously identified [26]. Our analysis with the new v4 pipeline and Gibbs sampling also revealed 49 DE transcripts associated with KEGG cancer and non-small cancer pathways that were not reported under the v3 pipeline with bootstrap sampling. Of particular interest was the detection of DE for the protein-coding transcripts *BCL2-201* (−5.83 log fold-change, adjusted p-value 7.105 × 10^−6^) and *BBC3-202* (1.33 log fold-change, adjusted p-value 6.164 × 10^−3^; Figure 2b). These results highlight the improvement in statistical power resulting from the latest analysis pipeline with *edgeR*-v4.

The overall agreement of DTE results was high among different pipelines. Yet, we observed a substantial number of DE transcripts that were exclusively detected in analyses using Gibbs sampling (n = 1,273; Figure 5c). This was a result of the higher resolution and lower RTA-dispersions provided by Gibbs sampling for lowly abundant transcripts and the consequent inclusion of such transcripts in the statistical analysis.

Specifically, a set of 2,355 transcripts were filtered out with *filterByExpr* exclusively in analyses using bootstrap sampling. Although transcripts exclusively reported as DE under the Gibbs sampling pipeline were, in general, more lowly expressed (median average log CPM -0.55; Supplementary Figure S10), several of them were associated with cancer and non-small cancer pathways (Figure 5d). This was the case for transcript *BAX-212* from the apoptosis regulator gene *BAX* (1.66 log fold-change, adjusted p-value 8.784 × 10^−4^). The improvement in computing time and the increase in power to detect DE transcripts between the human NCI-H1975 and HCC827 cell lines were in line with our previous observations based on our simulation study, which highlight *Salmon* with Gibbs sampling followed by *edgeR*-v4 as the pipeline of choice for transcript-level DE analyses.

## Discussion

Here, we present a faster and more accurate pipeline for differential analysis of RNA-seq data at the transcript-level with *edgeR* v4 and *Salmon*’s Gibbs sampling algorithm. Our simulation study demonstrates that the *edgeR*-v4 implementation provides uniformly more powerful DTE analyses than other methods. The bias-corrected QL method from *edgeR*-v4 allows the estimation of transcript-wise NB-dispersions to be bypassed, making our DTE workflow the fastest available. Count scaling can be straightforwardly applied in *edgeR* with estimated Gibbs-based RTA-dispersions to leverage the high computational speed of *Salmon*’s Gibbs sampling algorithm as well as its high resolution for lowly abundant transcripts.

For small-scale RNA-seq experiments, we expect our proposed pipeline to provide end-to-end DTE analyses in no more than half of the total computing time that was previously reported with bootstrap sampling by Baldoni *et al*. [13], with greater reductions for larger datasets. Applications of our recommended pipeline to the human adenocarcinoma cell lines dataset resulted in faster and more powerful analyses with several DE transcripts, including novel transcripts associated to key cancer-related genes that were not previously detected with *Salmon*’s bootstrap sampling.

The degrees of freedom available for estimating the RTA-dispersions is *N* (*B* − 1) where *N* is the total number of RNA samples and *B* is the number of technical samples per RNA sample. We found that the accuracy of the DTE results was not very sensitive to the number of Gibbs or bootstrap samples generated, and that fewer technical replicates are required for good accuracy than is usual for either *Salmon* or *kallisto*. We recommend that the number of Gibbs samples per RNA-seq library, *B*, should be chosen so that the *N* (*B* − 1) is at least 60 and preferably 200–300. Such a choice should be sufficient for optimal statistical power during DE assessment using *edgeR*-v4. For very large studies with *N* ≥ 200, just two resamples per RNA-seq library is sufficient for optimal results, making large-scale analyses very practical. If bootstrapping is used instead of Gibbs, then optimal results are already achieved with *N* (*B* − 1) about 50, with little further gain from drawing further technical samples. The optimal accuracy achievable by bootstrapping, however, is generally lower than that for Gibbs sampling, especially for larger experiments with more RNA samples.

Our simulation study assumed the same RNA-seq setup that we have previously recommended for DTE studies, with paired-end sequencing and at least 50 million 100bp read-pairs per sample [13]. Studies using single-end reads or shorter sequence reads would expect to observe somewhat larger RTA-dispersions on average, but the number of Gibbs or bootstrap samples required for optimal power should remain much the same.

As in our previous studies [6, 13], the main simulations were set to emulate small-scale designed experiments using model organisms, such as genetically identical mice or cell lines, with relatively low levels of biological variation. Simulations were also generated with more samples and a higher level of biological variability, in order to emulate a large-scale observational RNA-seq study with human subjects. *edgeR*-v4 with *Salmon* quantification and Gibbs sampling was consistently the most accurate pipeline across all simulations. More generally, *edgeR*-v3 and *edgeR*-v4 were the only analysis pipelines to control the FDR correctly for all sample sizes, with *edgeR*-v3 slightly more conservative than *edgeR*-v4.

This article has focused on pipelines using transcript quantification from *Salmon* or *kallisto*, because of their speed and popularity, but the *edgeR* DTE analysis pipeline examined here can also be used with RSEM output [8] via *edgeR*’s *catchRSEM* function.

## Code availability

The *catchSalmon* function is available in the *edgeR* Bioconductor package at https://bioconductor.org/packages/edgeR. The function imports fractional transcript counts and Gibbs resamples from *Salmon* to estimate the transcript-specific RTA-dispersion resulting from the RNA-seq quantification step. When performing DTE analyses with *edgeR*, users should divide transcript-level RNA-seq counts by the associated RTA-dispersion estimates. The function *glmQLFit* implements the latest bias-corrected QL method with adjusted deviances. Data and code to reproduce the results presented in this article are available at https://github.com/plbaldoni/GibbsDTE-code.

The versions of software used in the paper are: *ComplexHeatmap*: 2.18.0 [27], *edgeR*: 4.0.16, *fishpond* (*Swish* method): 2.8.0, *Gviz*: 1.46.1 [28], *limma*: 3.58.1, *R*: 4.3.2, *Rsubread* : 2.16.1, *sleuth*: 0.30.0, *Salmon*: 1.10.2, *wasabi* : 1.0.1.

## Data availability

The RNA-seq experiments analyzed here are available from the NCBI Gene Expression Omnibus with the accession numbers GSE60450 and GSE172421.

## Supplementary data

Supplementary Data are available in the file supp.pdf.

## Acknowledgments

The authors are grateful to Rob Patro for useful discussions.

