## Supplementary Material for "Faster and more accurate assessment of differential transcript expression with Gibbs sampling and edgeR v4"

6 October 2024

#### Contents

**1 Supplementary Figures** **2**

**2 Supplementary Table** **14**

[1]Bioinformatics Division, Walter and Eliza Hall Institute of Medical Research, Parkville, VIC 3052, Australia
[2]Department of Medical Biology, The University of Melbourne, Parkville, VIC 3010, Australia
[3]School of Mathematics and Statistics, The University of Melbourne, Parkville, VIC 3010, Australia

### 1 Supplementary Figures

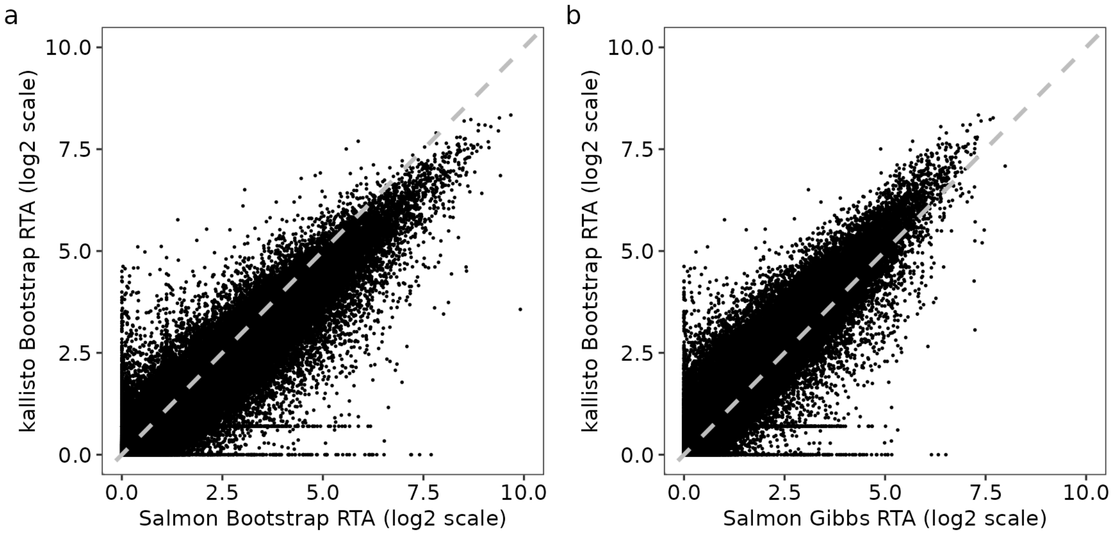


Figure S1: Transcript-level RTA-dispersion estimates from Salmon, obtained with either bootstrap or Gibbs sampling, and kallisto with bootstrap for the lung adenocarcinoma cell line data. (a) scatterplot of kallisto bootstrap- vs Salmon bootstrap-based RTA-dispersion estimates (log2 scale). (b) scatterplot of kallisto bootstrap- vs Salmon Gibbs-based RTA-dispersion estimates (log2 scale). RTA-dispersion estimates were obtained with 100 resamples. The line of equality is shown as a gray dashed line. All 137,788 expressed transcripts are shown.

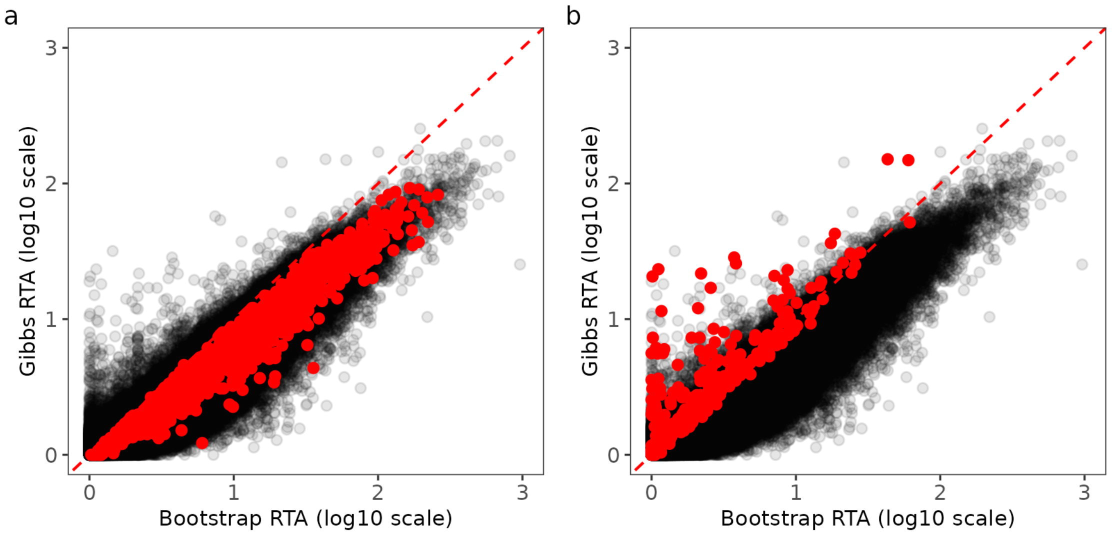


Figure S2: Transcript-level RTA overdispersion estimates for the lung adenocarcinoma cell line data. Both bootstrap- and Gibbs-based RTA estimates were obtained from *Salmon* quantification with 100 resamples. In panels (a) and (b), scatterplots showing Gibbs- and bootstrap-based RTA overdispersion estimates (log2 scale). In panel (a), red points highlight transcripts found to be DE between NCI-H1975 and HCC827 cell lines exclusively with Gibbs sampling. In panel (b), red points highlight transcripts found to be DE between NCI-H1975 and HCC827 cell lines exclusively with bootstrap sampling. Nominal FDR of 0.05 in analyses with *edgeR*-v4. All 137,788 expressed transcripts are shown.

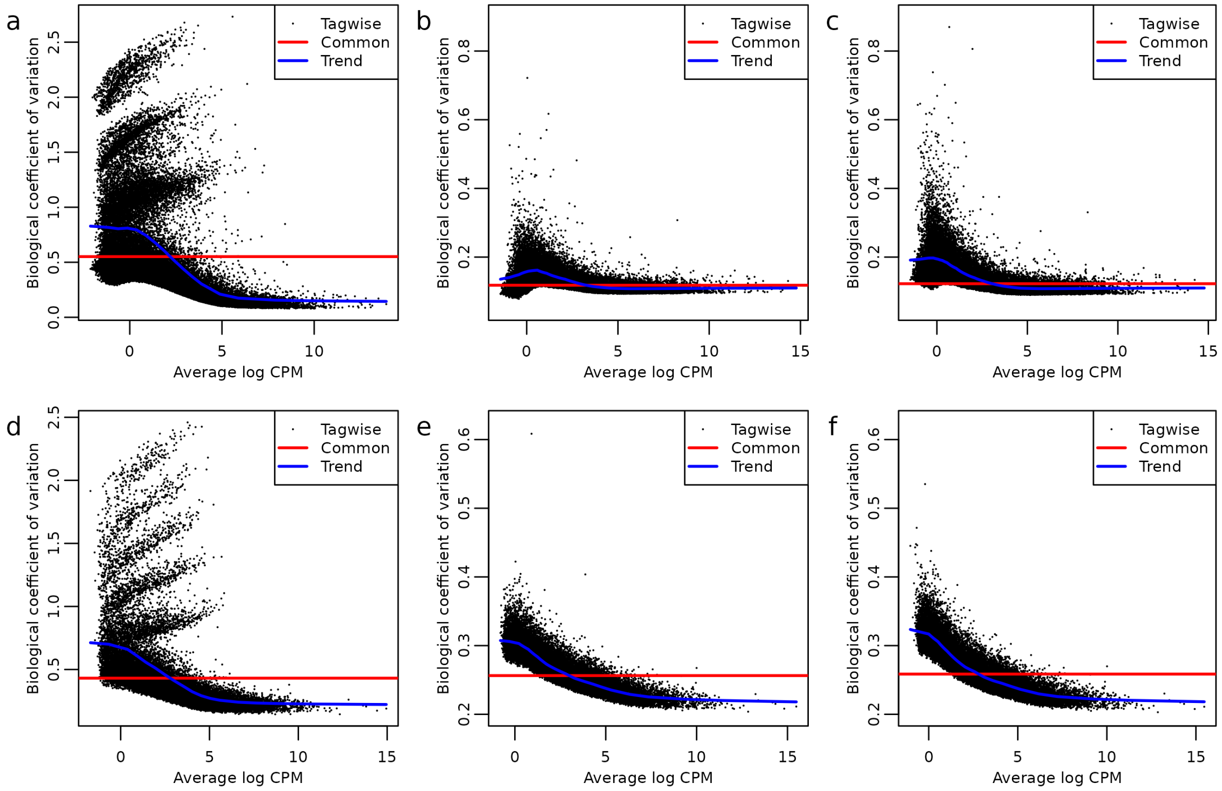


Figure S3: BCV plots of transcript-level counts before and after scaling for real and simulated RNA-seq data. Top panels (a–c) show results for the lung adenocarcinoma RNA-seq data with 80bp paired-end reads. Bottoms panels (d–f) show results from a simulated mouse RNA-seq experiment with 100bp paired-end reads, unbalanced library sizes, and five samples per group. Left panels (a) and (d) show BCV plots using original counts without scaling. Middle panels (b) and (e) show BCV plots after count scaling with bootstrap sampling. Right panels (c) and (f) show BCV plots after count scaling with Gibbs sampling. The plots without scaling show large NB dispersions that do not decrease with average expression. After scaling, the NB dispersions are much smaller and less variable and show a decreasing trend. Experiments were quantified with *Salmon* with 100 resamples.

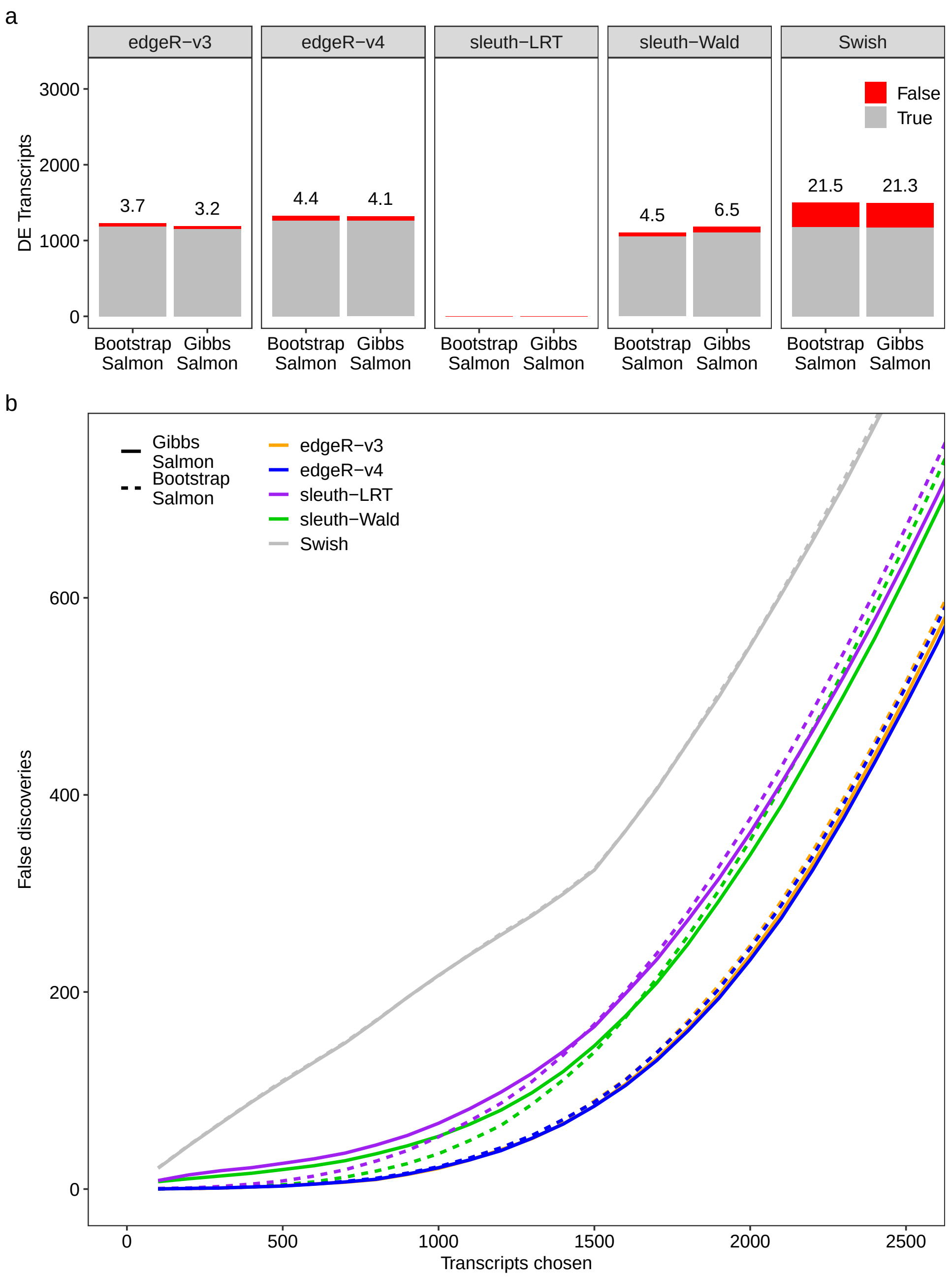


Figure S4: Simulation results from data generated with 100bp paired-end reads, unbalanced library sizes, and three samples per group. In panel (a), stacked barplots show the average number of true (gray) and false (red) positive DE transcripts at nominal 5% FDR for different DTE and sampling methods. The observed FDR is shown as a percentage over each bar. In panel (b), false discovery curves show the average number of false discoveries as a function of the number of chosen transcripts for different DTE and sampling methods. Transcript quantification was performed with *Salmon* with 100 bootstrap or Gibbs resamples. Results are averaged over 20 simulations.

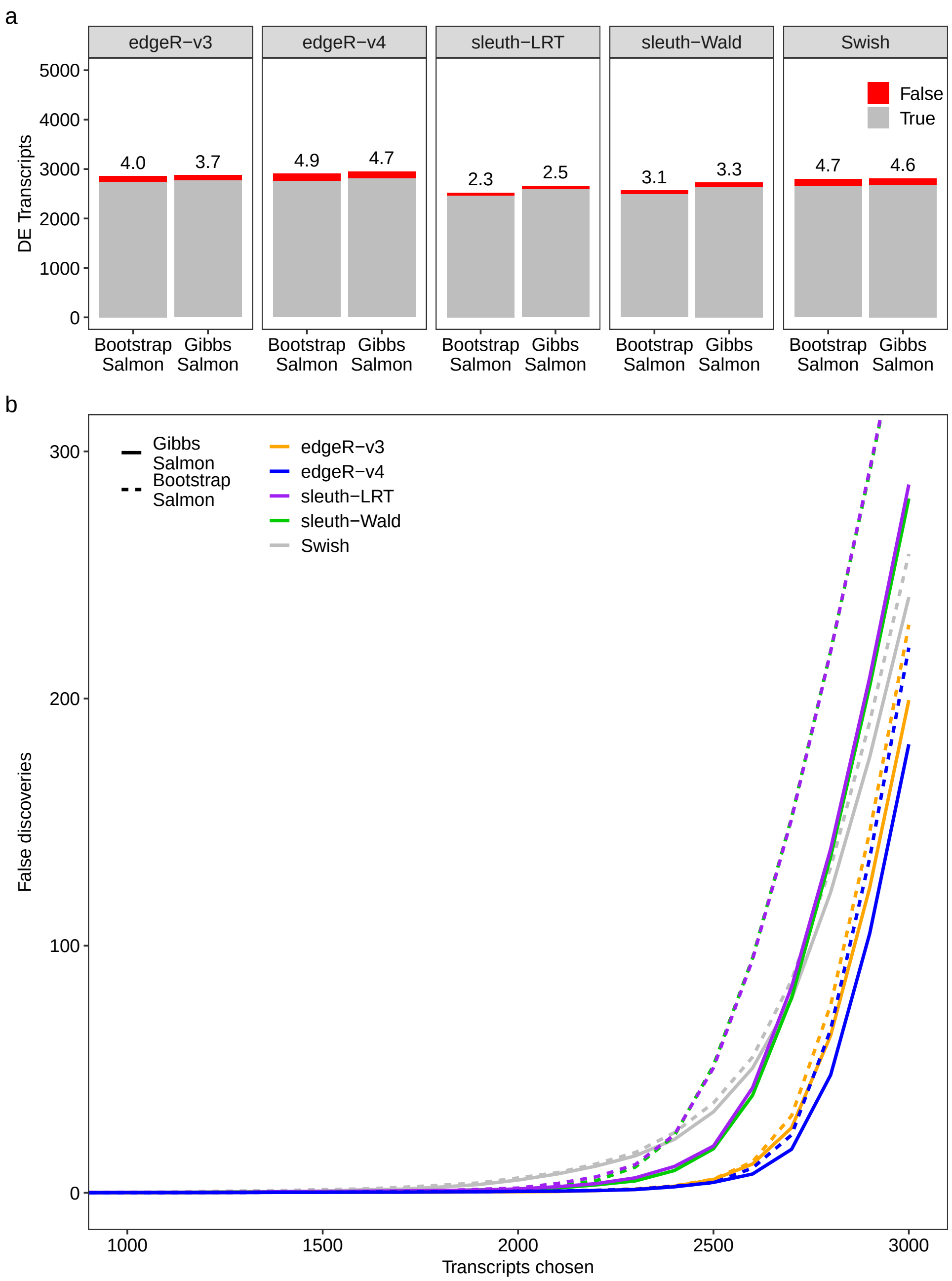


Figure S5: Simulation results from data generated with 100bp paired-end reads, unbalanced library sizes, and ten samples per group. In panel (a), stacked barplots show the average number of true (gray) and false (red) positive DE transcripts at nominal 5% FDR for different DTE and sampling methods. The observed FDR is shown as a percentage over each bar. In panel (b), false discovery curves show the average number of false discoveries as a function of the number of chosen transcripts for different DTE and sampling methods. Transcript quantification was performed with *Salmon* with 100 bootstrap or Gibbs resamples. Results are averaged over 20 simulations.

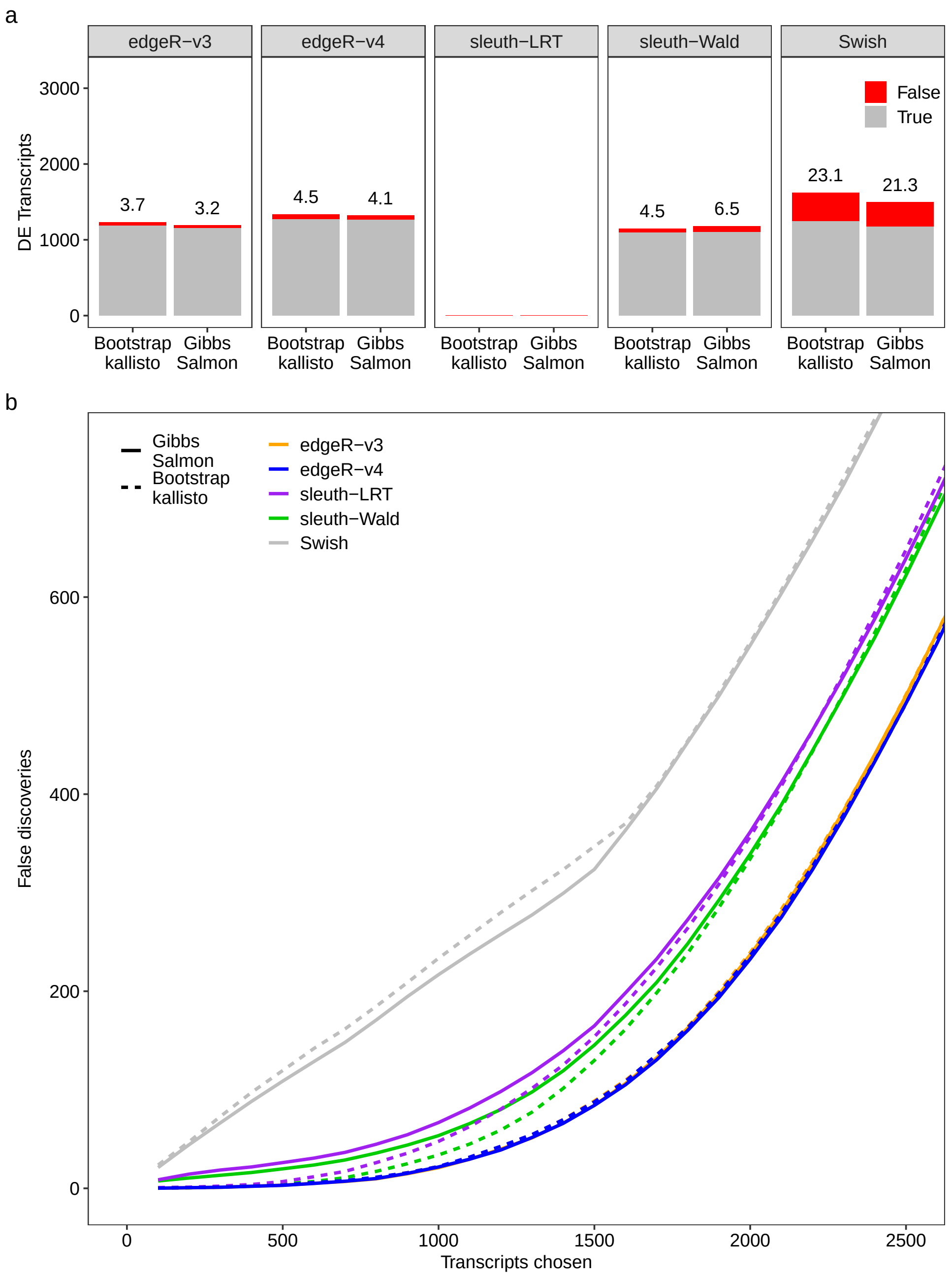


Figure S6: Simulation results from data generated with 100bp paired-end reads, unbalanced library sizes, and three samples per group. In panel (a), stacked barplots show the average number of true (gray) and false (red) positive DE transcripts at nominal 5% FDR for different DTE and sampling methods. The observed FDR is shown as a percentage over each bar. In panel (b), false discovery curves show the average number of false discoveries as a function of the number of chosen transcripts for different DTE and sampling methods. Transcript quantification was performed with *Salmon* (100 Gibbs resamples) and *kallisto* (100 bootstrap resamples). Results are averaged over 20 simulations.

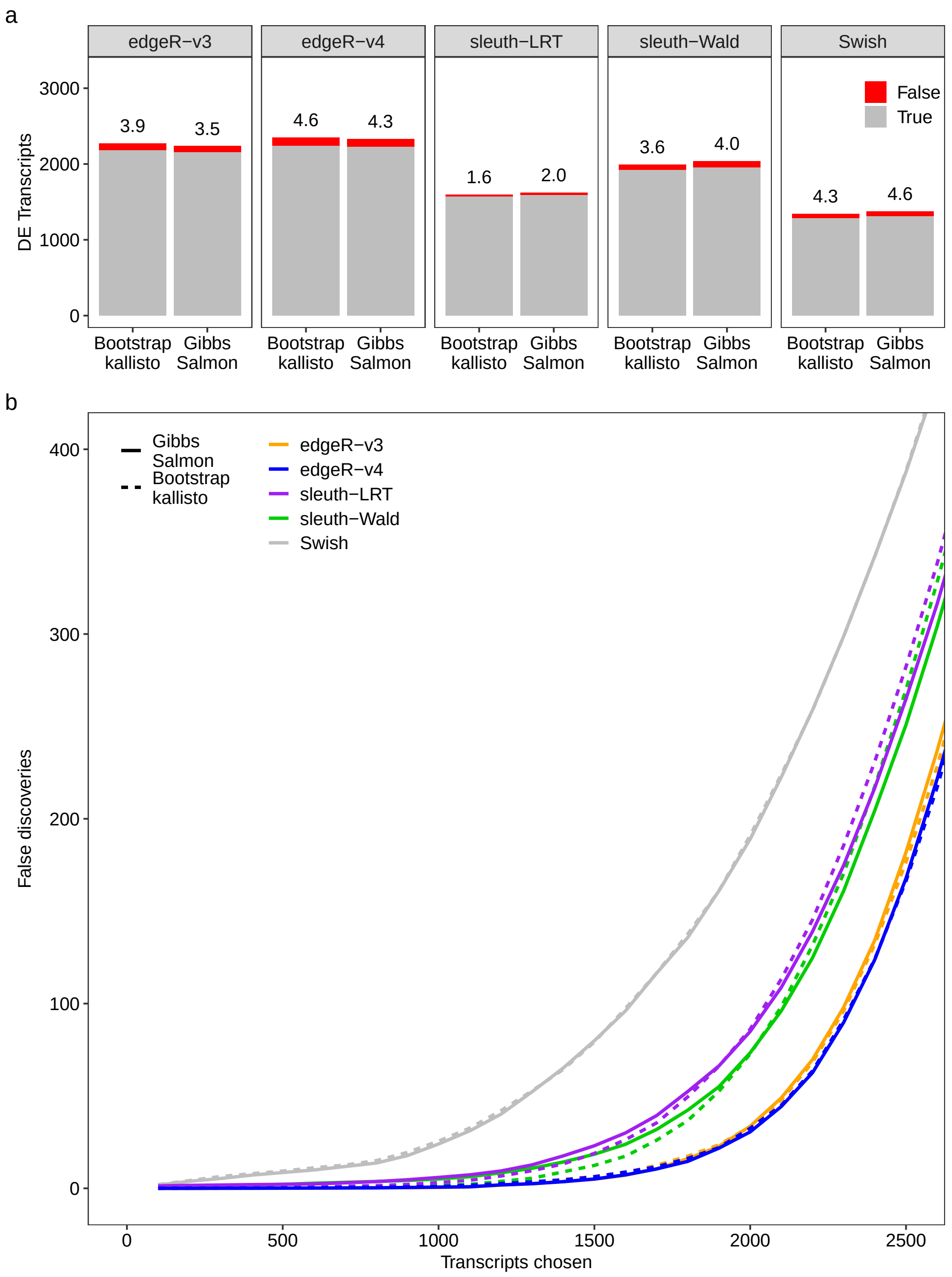


Figure S7: Simulation results from data generated with 100bp paired-end reads, unbalanced library sizes, and five samples per group. In panel (a), stacked barplots show the average number of true (gray) and false (red) positive DE transcripts at nominal 5% FDR for different DTE and sampling methods. The observed FDR is shown as a percentage over each bar. In panel (b), false discovery curves show the average number of false discoveries as a function of the number of chosen transcripts for different DTE and sampling methods. Transcript quantification was performed with *Salmon* (100 Gibbs resamples) and *kallisto* (100 bootstrap resamples). Results are averaged over 20 simulations.

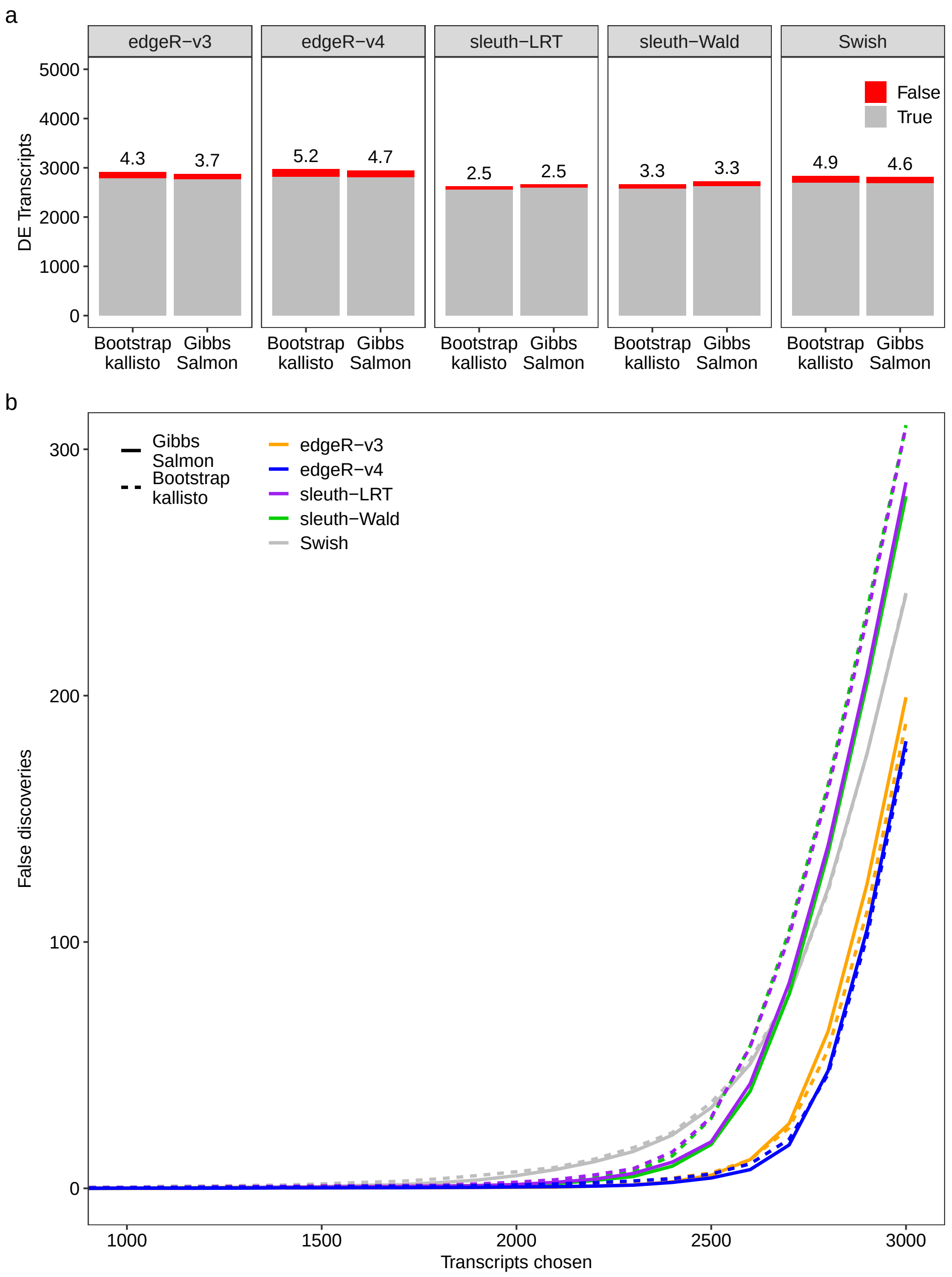


Figure S8: Simulation results from data generated with 100bp paired-end reads, unbalanced library sizes, and ten samples per group. In panel (a), stacked barplots show the average number of true (gray) and false (red) positive DE transcripts at nominal 5% FDR for different DTE and sampling methods. The observed FDR is shown as a percentage over each bar. In panel (b), false discovery curves show the average number of false discoveries as a function of the number of chosen transcripts for different DTE and sampling methods. Transcript quantification was performed with *Salmon* (100 Gibbs resamples) and *kallisto* (100 bootstrap resamples). Results are averaged over 20 simulations.

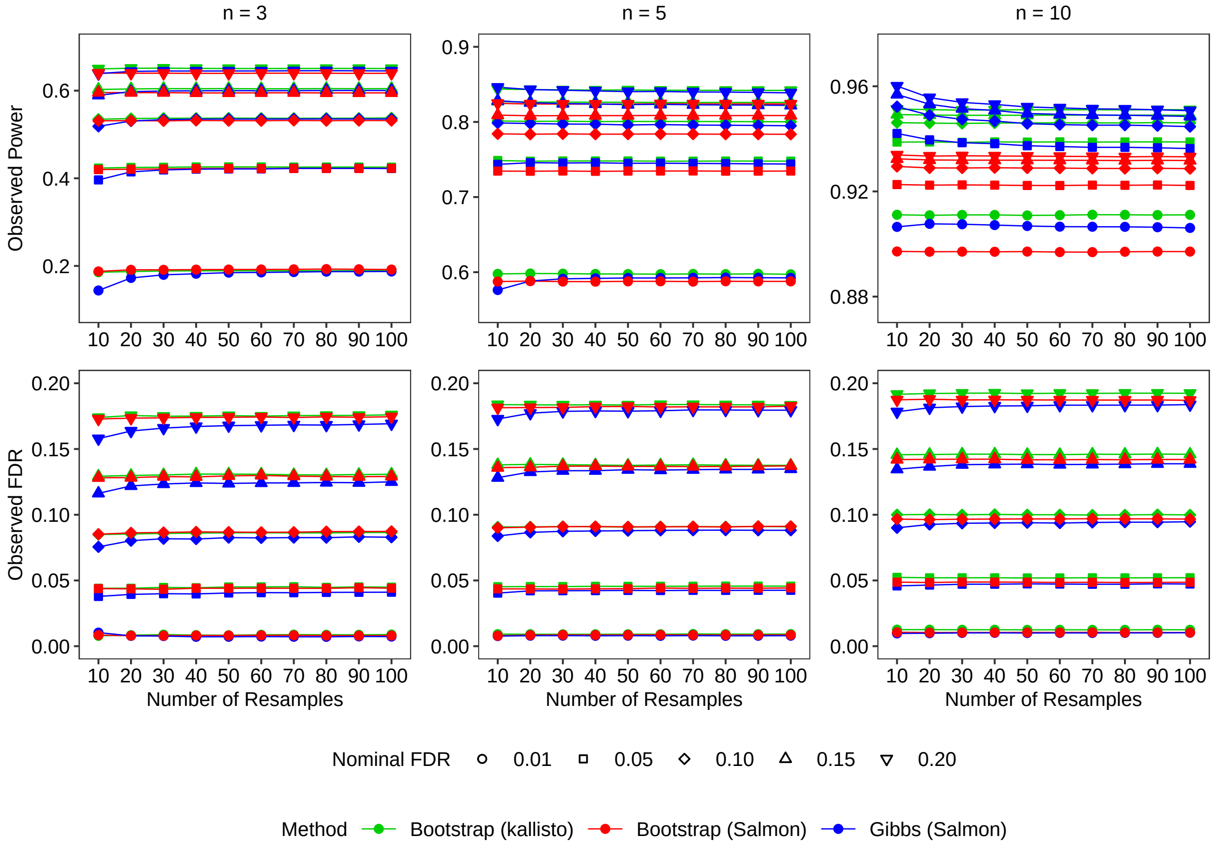


Figure S9: Plots showing the observed power and the observed false discovery rate (FDR) versus the number of replicate samples while controlling for different nominal FDR levels. Datasets were generated for two groups with $n = 3$, $n = 5$ or $n = 10$ samples per group. Results are presented for different number of samples per group and for Gibbs- and bootstrap-based quantification pipelines with *Salmon* and *kallisto*. DTE analyses were conducted with *edgeR*-v4. Simulated data generated with 100bp paired-end reads and unbalanced library sizes. Results are averaged over 20 simulations.

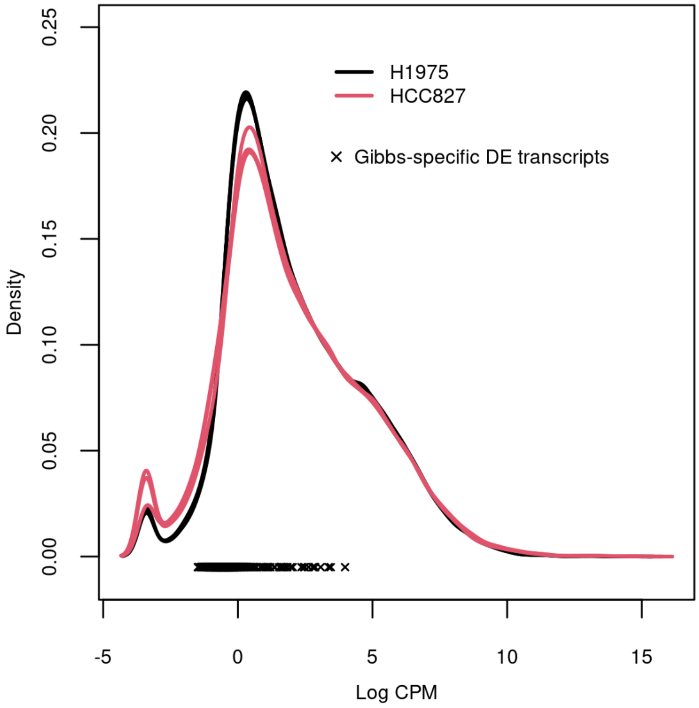


Figure S10: Plot showing density of log counts-per-million (CPM) from each of the RNA-seq samples of the human lung adenocarcinoma cell lines dataset. Crosses highlight transcripts found to be DE between NCI-H1975 and HCC827 cell lines exclusively with Gibbs sampling. Nominal FDR of 0.05 in DTE analyses.

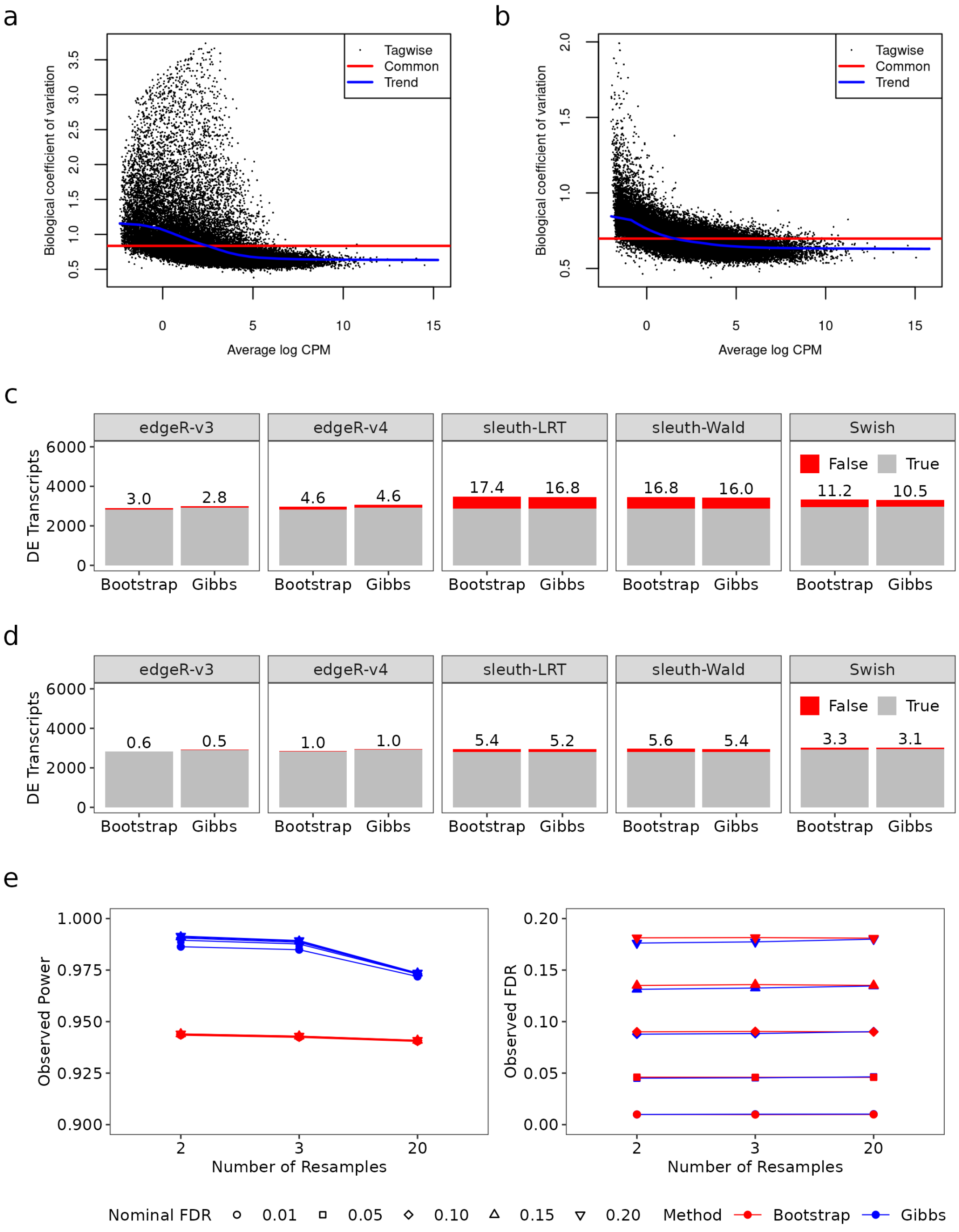


Figure S11: Simulation results from data generated with 100bp paired-end reads, unbalanced library sizes, and a hundred samples per group. In panels (a) and (b), BCV plots of transcript-level counts from a simulated dataset before and after count scaling, respectively. In panel (c), stacked barplots show the average number of true (gray) and false (red) positive DE transcripts at nominal 5% FDR for different DTE and sampling methods. In panel (d), stacked barplots show the average number of true (gray) and false (red) positive DE transcripts at nominal 1% FDR for different DTE and sampling methods. The observed FDR is shown as a percentage over each bar. In panel (e), plots showing the observed power and the observed false discovery rate (FDR) versus the number of replicate samples while controlling for different nominal FDR levels. Results are presented for both Gibbs- and bootstrap-based DTE pipelines with *Salmon* and *edgeR*-v4. Results are averaged over 20 simulations.

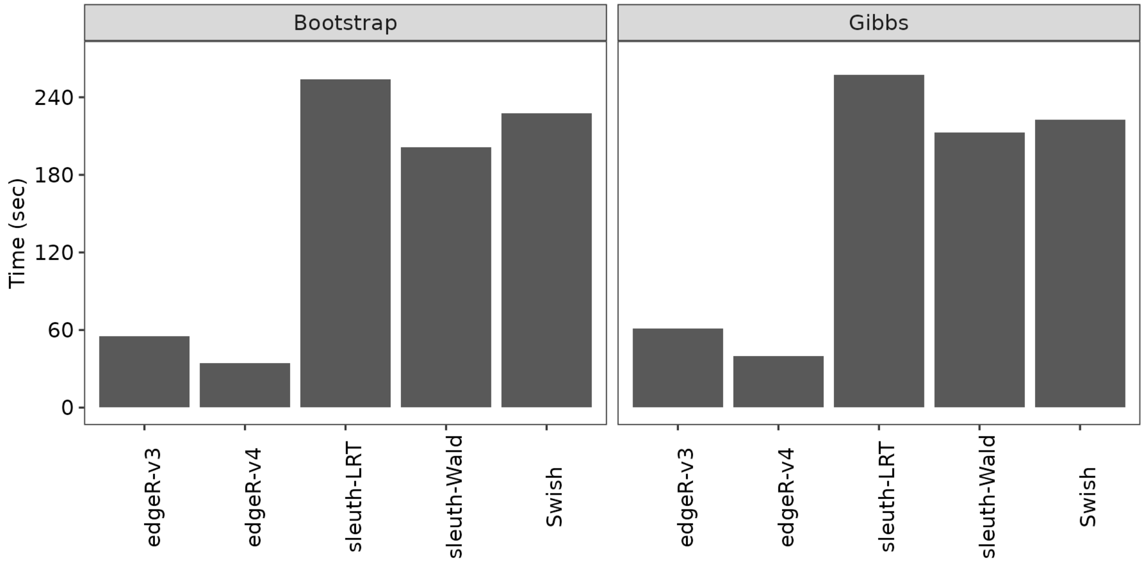


Figure S12: Computing time in seconds of DTE methods from data generated with 100bp paired-end reads, unbalanced library sizes, and a hundred samples per group. Results are presented for both Gibbs- and bootstrap-based DTE pipelines with *Salmon*. Results are averaged over 20 simulations.

### 2 Supplementary Table

Table S1: Total processing time per library with either Salmon (bootstrap or Gibbs sampling) or kallisto (bootstrap sampling). For *Salmon*, the total processing time is further split into the time spent during quantification, including algorithm initialization and optimization steps, and the time spent to generate the 100 bootstrap or Gibbs resamples. *kallisto* does not timestamp quantification and resampling time to log output. Results are averaged over all paired-end simulated libraries.

| Resampling Method | Library Size (million read pairs) | Quantification Time (minutes) | | |
|---|---|---|---|---|
| | | Optimization | Resampling | Total |
| Bootstrap (kallisto) | 25 | - | - | 4.92 |
| | 50 | - | - | 7.32 |
| | 100 | - | - | 11.71 |
| Bootstrap (Salmon) | 25 | 1.79 | 9.45 | 11.24 |
| | 50 | 3.22 | 13.41 | 16.64 |
| | 100 | 6.09 | 19.15 | 25.24 |
| Gibbs (Salmon) | 25 | 1.78 | 0.91 | 2.69 |
| | 50 | 3.22 | 1.62 | 4.85 |
| | 100 | 6.05 | 3.01 | 9.06 |
